# Tau Isoform Expression Drives Disease Outcomes Following a Single Closed Head Injury

**DOI:** 10.64898/2026.07.08.737276

**Authors:** Rachel Furhang, Riley Morrone, Elena Nikulina, Mahdavi Jere, Ashmeet Kaur, Fizza Nayab, Takashi Saito, Takaomi C. Saido, Peter J. Bergold

## Abstract

Tau protein has been implicated as an important mediator of traumatic brain injury (TBI). Adult human brain expresses 6 tau isoforms expressing 3 (3R) or 4 (4R) microtubule binding sites; adult mouse brain expresses only 4R tau. A role for tau isoform expression on TBI disease course is tested using wild-type C57/BL6 mice (WT) and C57/BL6 with a knocked-in human tau coding region (MAPTKI). Uninjured WT and MAPTKI mice have similar brain histology and behavior as they age. At subacute times (14 days post-injury (DPI)), injured MAPTKI mice have less white matter damage with similar neuronal loss as WT. At chronic times (90DPI), MAPTKI mice demyelinate while WT mice remyelinate. At 14DPI, tau phosphorylation differs between WT and MAPTKI mice. At 90DPI, thioflavin-S^+^ protein aggregates in MAPTKI corpus callosum are higher than WT. At 14 or 90DPI, WT and MAPTKI mice acquire Barnes maze, WT retention is impaired at 14DPI and MAPTKI retention impaired at 90DPI. At 14DPI, only MAPTKI mice acquire and retain active place avoidance; at 90DPI, only WT mice acquire active place avoidance. At 14DPI, only injured MAPTKI mice acquire alternating T-maze. These data suggest that WT and MAPTKI differ in both subacute and chronic disease course. At 14DPI, WT mice have greater white matter damage and behavioral impairments than MAPTKI mice. At 90DPI, impairments in WT mice partially recover, yet worsen in MAPTKI mice. This data suggests that 3R tau isoform expression alters the disease course of head injury.

**Highlights:** Post-injury disease course of MAPTKI mice expressing 3R and 4R tau differs from wild-type mice expressing only 4R tau.

At subacute times post-injury, MAPTKI mice have less white matter, yet similar gray matter, injury than wild-type mice.

At chronic times post-injury, white matter damage in MAPTKI worsens.

At subacute times post-injury, MAPTKI mice have fewer behavioral deficits than wild type mice. At chronic times post-injury, MAPTKI mice develop behavioral deficits not present at subacute times.

## Introduction

TBI is a major health issue with 2.35 million new cases of TBI in the United States in 2016 [1]. Chronic disabilities due to TBI affects 1 to 2 percent of the United States population [2, 3]. TBI severity ranges from mild to severe. Chronic impairments following TBI can occur at any injury severity but is more common as TBI severity increases [3]. TBI induces chronic cognitive deficits [4, 5], emotional regulation problems [6], personality changes [7] as well as neurodegeneration [2], epilepsy [8], and neuroendocrine disorders [9].

Despite the importance of the chronic consequences of head injury, few preclinical studies have assessed chronic long-term outcomes after a single TBI [10]. Memory deficits in male mice have been reported up to two months after injury [11–14]. It remains uncertain if mice ultimately recover from these memory deficits. In addition, these preclinical studies did not assess female mice.

Clinical studies suggest that females recover from cognitive and social deficits better than males [15]. Injured females, however, exhibit more anxiety and ICD-10-CM post-concussive symptoms [15]. Preclinical TBI studies assessing acute and subacute outcomes have not revealed a clear role for sex. Female mice have a decreased microglial response to TBI [16], yet lesion size lacks a sex effect [17]. Injured females perform better on motor tasks than males [18]. Cognitive tasks such as Morris water maze and Barnes maze yield conflicting results depending on the task, anesthetic used, and method of injury [18]. Generally, males are thought to have greater deficits than females [18].

This study injures mice using a closed-head model of TBI in which an electromagneic piston laterally strikes the head above the parietal cortex. The impact produces a rapid acceleraion-deceleraion with angular force and rotaional movement (Grin’kina et al., 2016). The impact also compresses the brain through the skull. The rapid head movement and brain compression results in transient apnea, loss of righing reflex as well as hemorrhage and contusion proximal to the impact site [19, 20]. The palern of injury immediately amer impact suggests that CHI models moderate TBI.

CHI induces a progressive neurodegeneraion as seen by widespread atrophy between 14 and 180 DPI [21]. Atrophy is accompanied by an increase in the density of phosphotau-immunoreactive (pTau+) cells in the ipsilesional cortex and thalamus, and in the ipsilesional and contralesional corpus callosum. Between 14 and 180 DPI, pTau^ββ^ cell density increases bilaterally in corpus callosum, yet decreases in cortex and thalamus [21]. DeepLabCut analysis of motor performance reveals late-onset motor deficits that persist at 180 DPI suggesting a chronic and progressive decline of motor function [22]. T2 and DT-MRI analysis show that CHI produces a diffuse axonal injury in brain regions both ipsilesional and contralesional to the impact site [21]. Injured WT mice are impaired on Barnes maze and active place avoidance at 14 DPI [20, 23]. It is uncertain if these cognitive deficits persist into more chronic times following CHI.

This study uses MAPTKI mice to examine whether human tau isoform expression alters subacute and chronic disease course following a single CHI. MAPTKI mice have an 80 mb knock-in of the human tau coding region under the control of the murine tau promotor [24]. The human or mouse tau locus expresses 6 isoforms containing 3 (3R) or 4 (4R) microtubule-binding domains [25]. Adult mice express only the three 4R tau isoforms. The additional microtubule binding site in 4R tau results in 3-fold stronger microtubule binding than 3R tau as well as more efficient microtubule assembly and less efficient microtubule shortening [26–31]. In contrast, 3R tau produces more flexible and dynamic microtubules [31].

Adult humans and MAPTKI mice express the three 3R and three 4R human tau isoforms while WT mice express only the three 4R tau isoforms [24, 32]. WT and MAPTKI mice express similar amounts of total tau protein. Between 6 and 24 months of age, MAPTKI and WT mice perform similarly on object location or novel object recognition tasks, radial arm water maze, and open field [32]. MAPTKI mice have fewer age-dependent motor deficits than WT mice. WT and MAPTKI brains have similar densities of astrocyte, oligodendrocyte, or microglia [32]. The hippocampus of WT and MAPTKI mice express similar levels of NeuN, MAP2 β-III tubulin, synapsin-1, and post-synaptic density protein-95 (PSD-95). The corpus callosum of both strains express similar levels of GFAP, Iba-1 and myelin basic protein [32]. Most importantly, in contrast to all other humanized tau mouse models, uninjured MAPTKI mice do not develop age-dependent tauopathy nor neurodegeneration when assessed up to two years of age.

Tau phosphorylation regulates microtubule binding [33]. Hyperphosphorylation promotes conversion of tau to pathological forms [34]. Increasing tau phosphate content by 3-fold decreases its solubility and promotes aggregation [35]. This study uses three antibodies (AT8, S214, and PHF-1) that recognize unique phosphorylation sites on the tau protein and do not differentiate between 3R and 4R tau [36–38].

The AT8 antibody assesses tau phosphorylation at Ser199, Ser202 and Thr205. Phosphorylation of these amino acids occur early in tau aggregation and are linked to disrupted axonal transport [39]. The PHF antibody recognizes phosphorylation at Ser396 and Ser404 [37]. Ser396 and Ser404 phosphorylation also inhibits microtubule binding and promotes aggregate formation [40]. Phosphorylation at Ser214 inhibits microtubule binding and promotes early tau aggregation [41]. Ser214 phosphorylation also alters synaptic function by preventing tau interaction with fyn kinase [42]. Thioflavin-S staining binds to β -sheet structures present in pathological protein aggregates and measures aggregate formation [43, 44]. Between 14 and 180 DPI, injured WT mice increase pTau^+^ cells, bilaterally in corpus callosum, but decreases in cortex and thalamus [21]. At 180 DPI, the injured ipsilesional WT corpus callosum increases tau phosphorylation as assessed by S214, PHF1 and AT8. Therefore, the same antibodies assess tau phosphorylation in injured WT and MAPTKI mice at 14 and 90 DPI.

This study assesses subacute and chronic cognitive deficits using Barnes maze, active place avoidance and alternating T-maze. In Barnes maze, mice learn an escape hole location based upon landmarks positioned outside the arena [45]. Acquiring Barnes maze activates many connected brain structures, including hippocampus, thalamus, and cortex [46]. Active place avoidance tests the ability to avoid a stationary shock zone on a rotating circular arena. Learning the shock zone location requires the mouse to attend to distal cues while ignore local olfactory cues that rotate with the arena [47–49]. Active place avoidance has higher cognitive demand than purely spatial tasks such as Barnes maze since the mouse learns the shock zone location by segregating relevant and irrelevant cues [48, 49]. Acquisition of active place avoidance is impaired by lesions to one hippocampus, bilateral lesions of the retrosplenial cortex or local demyelination of the fimbria fornix [47, 50, 51]. The chronic persistence of these deficits is unknown. Alternating T-maze tests working memory using the innate tendency of mice to explore novel environments [52]. Alternating T-maze task is highly sensitive to lesions of the hippocampus and thalamus [53, 54].

In this study, WT and MAPTKI receive similar initial injury, but have differing outcomes at both subacute and chronic times post-injury. This suggests a key role for tau isoform expression in disease course following a single CHI.

### Animals

A colony of MAPTKI mice (B6.Cg-Mapt<tm1.1(MAPT)Tcs) was generated from frozen sperm provided by Dr. Takaomi C. Saido, Riken BioResource Research Center (Tsukuba, Japan). and homozygote crosses of male and female MAPTKI mice [24, 55]. MAPTKI mice are genotyped as described by Saito et al. [24]. At every fifth generation, homozygous MAPTKI mice are back-crossed with a WT C57Bl/6J mouse. After weaning, each MAPTKI mouse receives a unique identification number. A total of 212 mice (149 males, 133 females) are used. Twenty-two injured mice are excluded due to a return of righting reflex greater than 450 seconds. Upon arrival to SUNY-Downstate, male and female C57Bl/6J mice (28-30g, Jackson Laboratories) (WT) receive a unique identification number. Both WT and MAPTKI mice are randomized to receive either sham-injury or closed head injury (CHI). MAPTKI litters are evenly distributed by sex to sham or injury groups. Each group contains MAPTKI mice from multiple litters.

### Closed head injury

Closed head injury is done as described by Lawless, et al [22]. In the CHI murine TBI model, the mouse head is struck above the parietal cortex with an electromagnetic piston. Impact produces an acceleration-deceleration of the head resulting in angular and rotational movements as well as compression of the brain through the skull [19]. This produces contusion and hemorrhage at the impact site with distal gray and white matter damage [19–21]. Immediately prior to injury, deep anesthesia is induced (3% isoflurane in oxygen (2.0 L/min)) and maintained (2% isoflurane in oxygen (2.0 L/min)) until after impact. The shaved mouse head is put in a modified Kopf stereotaxic apparatus; the apparatus bed and ear bar holders are covered with 0.5 inches of polyurethane foam. An electromagnetic impactor (NeuroPactor™ NeuroscienceTools, O’Fallon, MO) with a 4.0 mm diameter impactor tip strikes the head at 6.4 m/s above the right hemisphere located 3 mm lateral from the mouse midline and 3 mm rostral from the anterior end of the ears. After impact, the mouse is placed on its side on a 37°C heated pad and the time recorded to resume an upright position. Sham mice receive the same procedure without an impact. Animals are returned to their home cage after regaining righting reflex. Time to regain righting reflex must be less than 120 seconds for inclusion in the sham group, and greater than 450 seconds for inclusion in the CHI group. Head movement during the CHI procedure results in an impact site that slightly varies (from Bregma, (in mm); AP, -0.47 - -0.83; DV, -1.8 - -2.2; ML 0.0 – 1.4). All animal studies are conducted at State University of New York (SUNY) Downstate Health Sciences University and approved by the Institutional Animal Care and Use Committee of SUNY Downstate (Animal welfare assurance A3260–01; Protocol 22-10631). All animal studies are performed in accordance with the NIH Guide for the Care and Use of Laboratory Animals and ARRIVE (Animal Research: Reporting of *In Vivo* Experiments) guidelines.

### Behavioral testing

All behavioral tests are done a windowless room with an average illumination of 268 lux and temperature of 24°C. At the end of each testing session, a 70% ethanol wash removes any residual odors on all behavioral apparatuses. Barnes maze is done as described by Whitney et al. starting at 7 or 83 days after sham-CHI or CHI [20]. The Barnes maze apparatus is a circular platform (92 cm) containing 20 equally spaced holes (5cm) located 1.5 meters from walls containing prominent visual landmarks. The escape box position is unchanged in all trials. Gently moving air from a fan located 270° from the escape hole provides additional mild motivation to find the escape hole. Barnes maze testing occurs over 5 days. On day 1, mice receive a 5-minute habituation trial. Five minutes after the habituation trial, mice receive four 3-minute acquisition trials with a 15-minute inter-trial interval. This regimen of acquisition trials is repeated on days 2-4. On day 5, mice receive a 90-second probe trial with the escape box removed. In all trials, the position of the mouse positions is tracked at 10 frames per second with a webcam (Tecknet C016 720p HD) located 1.5 m above the maze tracks using Debut video software. Videos are analyzed by a blind evaluator using AnyMaze software (Stoelting). Task acquisition is assessed by the average time to reach the escape hole over 4 daily trials. Task retention in the probe trial is assessed by the percent time that the mouse is in the quadrant containing the escape hole.

Active place avoidance (APA) is done with modifications of the method of Burghardt, et al [56]. During APA, mice avoid a stationary shock zone located on a 40cm rotating arena. The arena rotates one revolution per minute and contains a computer-generated shock zone. An infrared Firewire camera located 1.2m above the arena. Track analysis software (Bio-Signal Group Corp., Brooklyn, NY) assess mouse location and shock zone entries. Mice are habituated for 10 minutes on the rotating arena with an inactive shock zone and total distance traveled and speed is assessed. Mice then receive 2 × 10-minute sessions on the rotating arena with an active shock zone with a 60-min intertrial interval. The following day, mice receive 3 × 10-minute sessions. Three days later, mice receive a 10-minute probe trial. Total distance traveled, and shock zone entrances are assessed on training and probe trials.

Alternating T-maze is performed on an apparatus (70 × 10cm) containing a start arm (30 × 10 cm) connected to two goal arms (30 × 10cm). The base of each arm has a guillotine door. Mice are habituated to the T-maze for 5 minutes. A mouse is placed in the start arm with the guillotine door down. The door is opened and the animal chooses a goal arm. Once a goal arm is chosen, the door to the goal arm is closed confining the mouse in the goal arm for 30 seconds. The mouse animal is then placed back in the start arm with the door down. The procedure of choosing a goal arm is repeated five times and discrimination index (the percent of correct alterations per five trials) is assessed.

### Histology

Mice are deeply anesthetized with isoflurane (3%) in oxygen (2 L/min) and perfused transcardially first with phosphate buffered saline (PBS) at 4°C followed by paraformaldehyde (4% (w/v)) in PBS at 4°C for 3 minutes at a flow rate of 7mL/min. The brain is isolated and post-fixed in paraformaldehyde (4% (w/v)) in PBS for 48 hours at 4°C and then transferred to PBS and stored at 4°C. Brain are embedded in paraffin, and 5µm or 8µm sections prepared (HistoWiz, Brooklyn, NY). Brain regions analyzed are (from Bregma (in mm), Cortex, AP, -1.0 - -0.54; DV, 0.5-1.2; ML, 0.8-1.8; Corpus callosum, AP, -1.0 - -0.65; DV, 1.0-1.2; ML, 0.8-1.8; CA1 stratum pyramidale, AP, -1.3- -2.8; DV, 4.2-4.9; ML, 0.36-1.8; CA3 pyramidale, AP, -1.0- -2.0; DV, 4.0-4.8; ML, 0.36-1.8; Thalamus, AP, -0.6- -3.0; DV, 2.0-3.6; ML, 0.36-1.8). Chromogenic staining of NeuN (10 mg/mL, Abcam, Ab104225), AT8 (0.4 mg/mL, Thermo-Fischer, MN1020), S214 (0.2 mg/mL, Thermo-Fischer, 44-742G) and PHF1 (1:500, gift of P. Davies) is done the Mount Sinai Brain Bank and Research CoRe using antibodies on a Ventana Benchmark XT or Ventana Discovery Ultra. Antigens are retrieved using antigen retrieval buffer (Tris/Borate/ EDTA buffer, pH 8.0–8.5, 950-224, Roche Diagnostics, Basel Switzerland). Primary antibodies are diluted in antibody dilution buffer (ADB250, Ventana Medical System Inc., Roche Diagnostics) and visualized with secondary antibodies linked to horseradish peroxidase using OptiView 3,3ʹ-Diaminobenzidine (DAB) Detection kit (760-700, Roche Diagnostics. Slides are counterstained with hematoxylin. The Mount Sinai Brain Bank and Research CoRe have previously determined reagents concentrations. Whole slide brightfield scanning is performed using an Aperio CS2 slide scanner with a 40X lens. Bielschowsky’s silver stain is done with 5 μm sections to provide for efficient penetration of the stain while allowing adequate representation of the tissue structure. Silver-stained sections were scanned with an Aperio CS2 scanner as previously described [19]. QuPath software (Version 0.6.0) was manually trained to identify silver-stained structures similar in size to axonal bulbs (3-6 μm in diameter). One section per animal was assessed. Axonal bulbs are assessed in the ipsilesional and contralesional corpus callosum. Silver stain intensity is normalized to the superior cerebellar peduncle (AP, -4.2- -5.1; DV, 2.2-2.9; ML, 0.36-1.80), a brain region not injured in the CHI model.

### Immunofluorescence

Slides are deparaffinized and antigens retrieved using 10mM Sodium Citrate, 0.05% Tween 20, pH 6.0) at 96 °C for 30 minutes. Slices are rinsed in PBS plus 0.05% (v/v) Tween 20 (PBS, (in mM) NaCl 137, KCl, 2.7 Na_2_HPO_4_, 10mM and KH_2_PO_4_ 1.8) and permeabilized with PBS plus 0.15% (v/v) Triton X-100. Slides are washed in PBS plus Tween 20 (0.05% v/v), and incubated overnight in blocking buffer (bovine serum albumin (1%) normal donkey serum (1.5%) and NaN_3_ (1%) in PBS plus 0.05% (v/v) Tween 20 at 4 °C. Sections are treated with either anti-APP (Invitrogen 51-2700 1:500) or SMI-34 (Biolegend 835503 1:500) antibodies overnight at 4°C. Anti-APP Antibody detects full-length APP (APP695, 751, 770), N-terminal truncated APP forms and C-terminal membrane-anchored APP but not Amyloid beta. Slices are incubated overnight in either AlexaFluor 488 (1 mg/mL, Thermo-Fischer, A21202) or AlexaFluor 568, 1 mg/mL, Thermo-Fischer, A10042) in blocking buffer at 4 °C. Slides are washed with PBST scanned the same as SA2.1. QuPath automated positive cell detection assesses APP puncta positive density or SMI-34 staining intensity in corpus callosum.

Fluoromyelin staining: Deparaffinized coronal sections (5µm) are twice rehydrated in PBS and then treated for 30min at room temperature with FluoroMyelin™ Red Fluorescent Myelin Stain (1:300, ThermoFisher F34652). The sections are washed in 3 times for 10 minutes in PBS, cover-slipped and scanned with a Zeiss Axio Observer LSM 800 microscope. Fluoromyelin staining intensity in the corpus callosum is assessed and normalized to the columns of the fornix (AP, 0.13- -2.15; DV, 0.6-3.0; ML, 0.1-1.0).

Thioflavin-S staining: Slides are stained as described by Sun et al., 2002 [57]. Background auto-fluorescence is removed by sequential incubation in 0.3% KMnO_4_ (Millipore, 223468), 1% KHSO_4_ (Millipore, P2522) and 1% (COOH)_2_ (Millipore, 241172), and 1% NaBH_4_ (Millipore, 213462).

Thioflavin S (0.05%, MedChemExpress, HY-D0972) in 50% ethanol is added for 5 min at room temperature and washed with 80% ethanol and three changes of distilled water for 2 minutes each. Slides were scanned and digitized with a Zeiss Axio Observer LSM 800 microscope.

## Results

Differences in body weight alters traumatic brain injury severity [58]. Immediately before sham-CHI or CHI, WT and MAPTKI mice have similar body weights (Supplementary Fig. 1A). Immediately after impact, acute injury severity is assessed by the time needed for a regain righting reflex [19]. Time to regain righting reflex does not differ between male or female injured WT and MAPTKI mice suggesting similar initial injury in WT and MAPTKI mice (Supplementary Fig. 1B).

Gray and white matter damage in WT and MAPTKI mice is assessed at 14 and 90 DPI. Ispsilesional CA1 and CA3 NeuN^+^ neuronal density is lower in injured male WT mice at 14 DPI, [87]. NeuN^+^ cells are therefore assessed in male and female WT or MAPTKI mice at 14DPI and 90DPI in the hippocampus, cortex, and thalamus (Figs. 1,2). No significant sex effects are observed; therefore, results are combined from males and females (Table 1).

**Figure 1.**
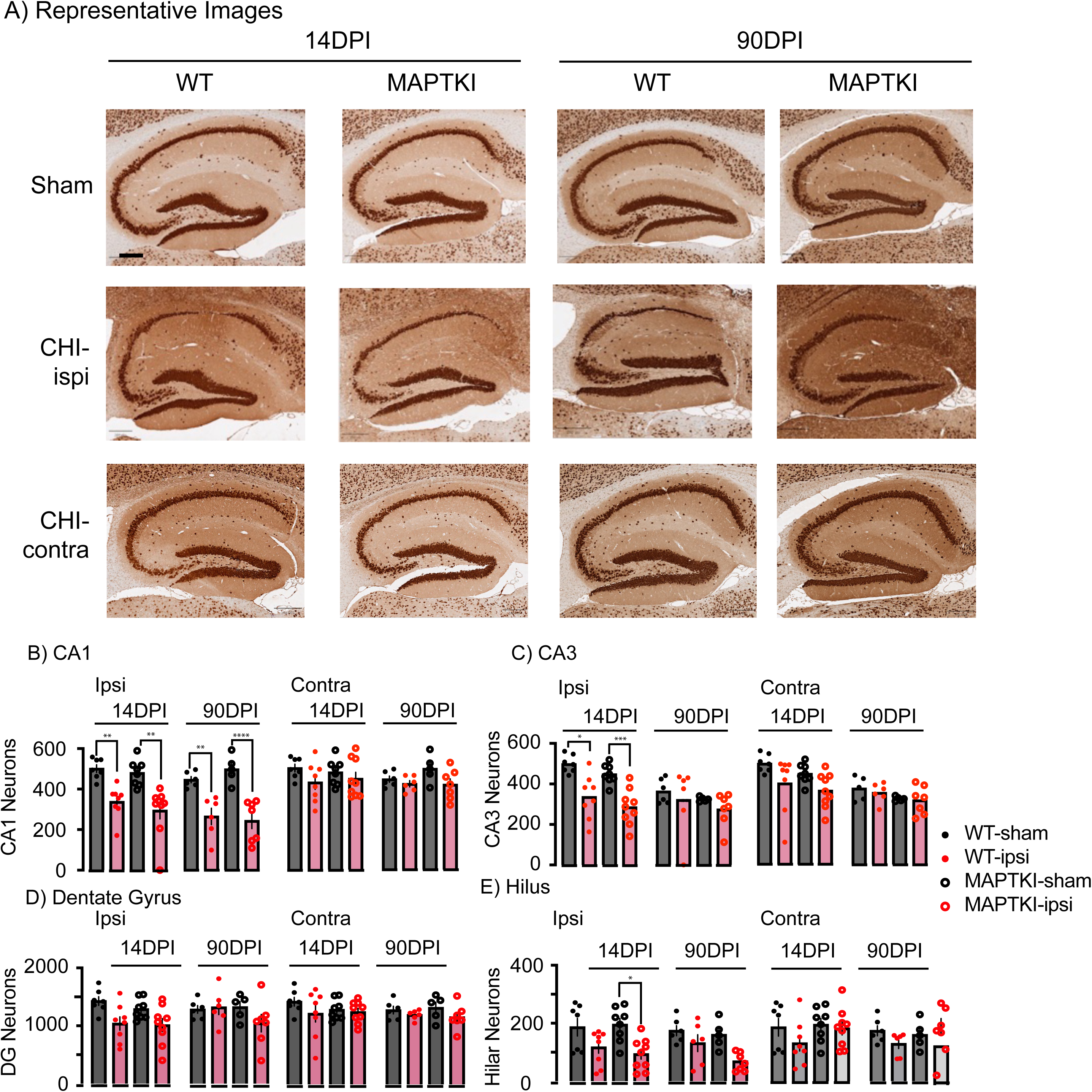
Similar Hippocampal neuronal loss in injured WT and MAPTKI mice. A) Representative images of the NeuN-stained ipsilesional and contralesional hippocampus. Scale bar 250µm. NeuN^+^ cell number is assessed in B) CA1, C) CA3, D) Dentate gyrus and E) Hilus. *p=0.02, **p=0.01, ***p=0.0009, ***p=0.0001. n’s=6-9.

**Figure 2.**
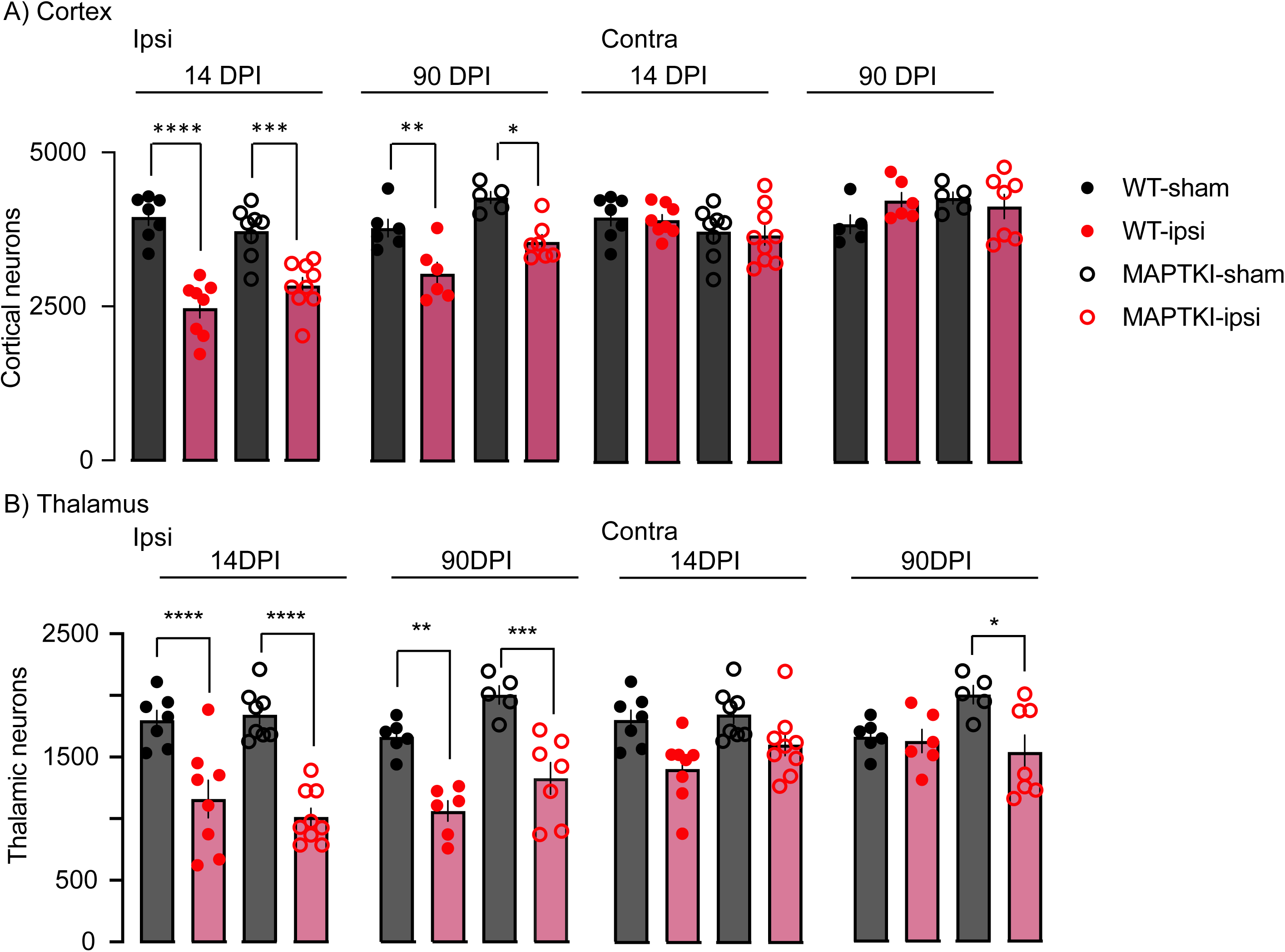
Similar cortical and thalamic neuronal loss in injured WT and MAPTKI mice. NeuN^+^ cell number is assessed in the ipsilesional and contralesional A) Cortex or B) Thalamus. *p=0.03, **p=0.005, ***p=0.001, ****p<0.0005. n’s=6-9.

**Table 1.** Two way ANOVA assessing neuronal density in hippocampus, cortex or thalamus for an effect of sex.

|  | CA1 WT |  | CA1 MAPTKI |  | CA3 WT |  | CA3 MAPTKI |  |
| --- | --- | --- | --- | --- | --- | --- | --- | --- |
|  | 14 DPI | 90 DPI | 14 DPI | 90 DPI | 14 DPI | 90 DPI | 14 DPI | 90 DPI |
| Sex | F<sub>1,10</sub>=11.9<br>4 p=0.006 | F<sub>1,8</sub>=3.0<br>p=0.1 | F<sub>1,13</sub>=6.9<br>p=0.02 | F<sub>1,8</sub>=15.9<br>p=0.004 | F<sub>1,10</sub>=8.0<br>p=0.02 | F<sub>1,8</sub>=2.3<br>p=0.2 | F<sub>1,13</sub>=20.1<br>p=0.0006 | F<sub>1,8</sub>=0.4<br>p=0.5 |
| If significant re<br>there biologically<br>relevant pairwise<br>comparisons? | No |  | No | No | No |  | No |  |
|  | DG WT |  | DG MAPTKI |  | Hilus WT |  | Hilus MAPTKI |  |
|  | 14 DPI | 90 DPI | 14 DPI | 90 DPI | 14 DPI | 90 DPI | 14 DPI | 90 DPI |
| Sex | F<sub>1,9</sub>=6.8<br>p=0.03 | F<sub>1,8</sub>=8.4<br>p=0.02 | F<sub>1,13</sub>=2.2<br>p=0.2 | F<sub>1,8</sub>=0.07<br>p=0.8 | F<sub>1,10</sub>=70.5<br>p<0.0001 | F<sub>1,8</sub>=3.2<br>p=0.1 | F<sub>1,13</sub>=0.1<br>p=0.7 | F<sub>1,8</sub>=4.1<br>p=0.07 |
| If significant, are<br>there biologically<br>relevant pairwise<br>comparisons? |  | No |  |  | No |  |  |  |
|  | Cortex WT |  | Cortex MAPTKI |  | Thalamus WT |  | Thalamus MAPTKI |  |
|  | 14 DPI | 90 DPI | 14 DPI | 90 DPI | 14 DPI | 90 DPI | 14 DPI | 90 DPI |
| Sex | F<sub>1,10</sub>=6.6<br>p=0.03 | F<sub>1,8</sub>=8.4<br>p=1.0 | F<sub>1,13</sub>=7.7<br>p=0.02 | F<sub>1,8</sub>=1.8<br>p=0.2 | F<sub>1,10</sub>=1.2<br>p=0.3 | F<sub>1,8</sub>=0.7<br>p=0.4 | F<sub>1,13</sub>=0.9<br>p=0.4 | F<sub>1,8</sub>=1.0<br>p=0.3 |
| If significant, are<br>there biologically<br>relevant pairwise<br>comparisons? | No |  | No |  |  |  |  |  |

NeuN^+^ cell number is assessed in hippocampal CA1, CA3, dentate gyrus and hilus (Fig. 1B-E, Table 1). Injury has a significant effect on ipsilesional CA1 with fewer WT or MAPTKI NeuN^+^ cells at 14 and 90DPI (Table 1, Fig. 1B). Contralesional CA1 also has a significant injury effect yet with no biologically significant pairwise comparisons (Table 1, Fig. 1B). Ipsilesional CA3 NeuN^+^ cell number has significant effects of time, injury, and time*injury with WT and MAPTKI mice having fewer NeuN^+^ cells at 14DPI (Table 1, Fig. 1C). Contralesional CA3 NeuN^+^ cell number has significant effects of time, genotype injury and time*genotype, yet with no significant pairwise comparisons (Table 1, Fig. 1C). NeuN^+^ cell number in the ipsilesional and contralesional dentate gyrus have significant injury effects yet with no significant pairwise comparisons (Table 2, Fig. 1D). NeuN^+^ cell number in the ipsilesional and contralesional hilus have significant injury effect, with the ipsilesional hilus of MAPTKI mice having fewer NeuN^+^ cells at 14DPI (Table 2, Fig. 1E). Ipsilesional cortical NeuN^+^ cell number has significant effects of time, genotype, injury, time*genotype and time*injury with WT and MAPTKI mice with fewer NeuN^+^ cells at both 14 and 90DPI (Table 3, Fig. 2A). In contrast, injury has no effect on contralesional cortical NeuN^+^ cell number. Both ipsilesional and contralesional thalamic NeuN^+^ cell number have significant effects of injury and injury*genotype (Table 3). At 14 and 90 DPI, injury lowers NeuN^+^ cells in ipsilesional thalamus of both strain (Fig. 2B). At 90 DPI, injured MAPTKI mice also have fewer contralesional thalamic NeuN^+^ cells (Fig. 2B). These results suggest similar hippocampal and cortical neuronal loss in MAPTKI and WT mice. In contrast, injured MAPTKI mice have greater thalamic neuronal loss.

**Table 2.**
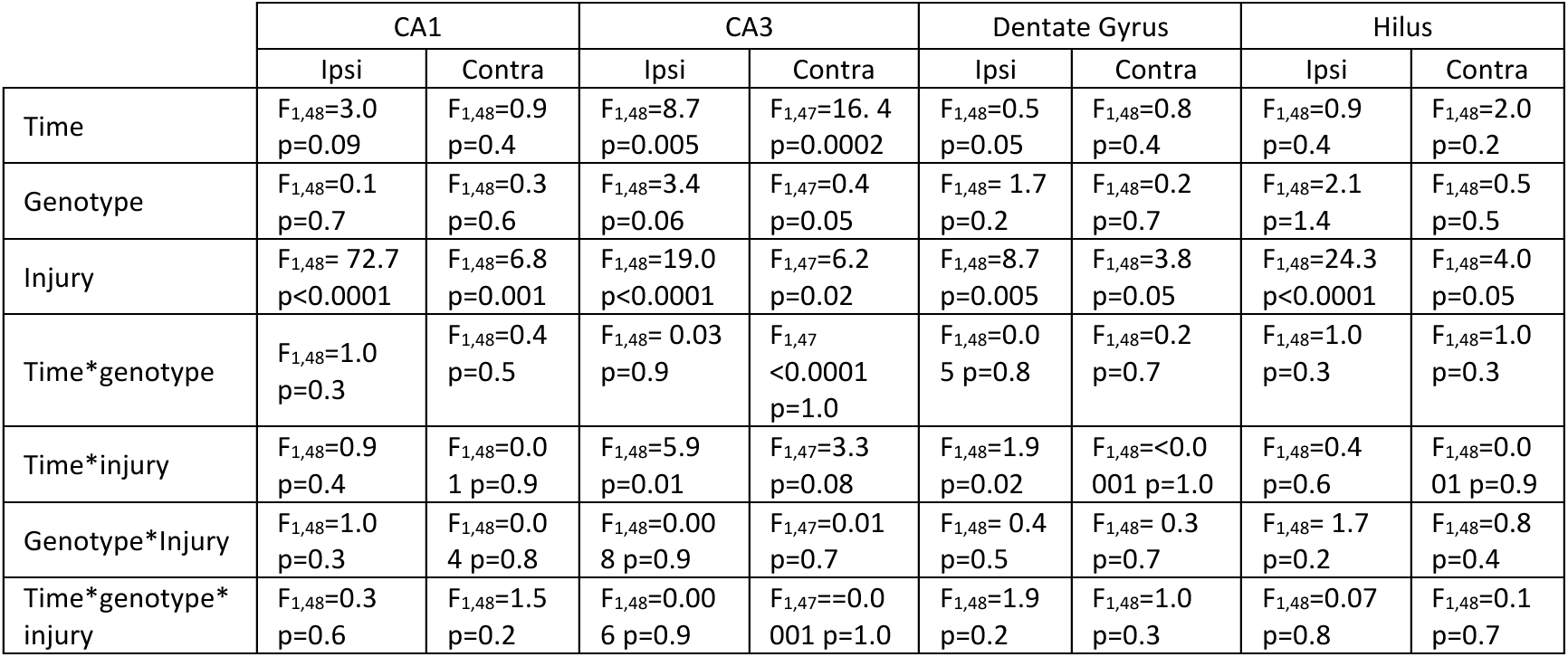
Three-Way ANOVA assessing hippocampal neuronal density for an effect of time, genotype or injury.

**Table 3.** Three-Way ANOVA assessing cortical and thalamic neuronal density for an effect of time, genotype or injury.

|  | Cortex |  | Thalamus |  |
| --- | --- | --- | --- | --- |
|  | Ipsi | Contra | Ipsi | Contra |
| Time | $F_{1,48}=15.8$ $p=0.0002$ | $F_{1,47}=8.5$ $p=0.006$ | $F_{1,48}=0.7$ $p=0.4$ | $F_{1,48}=0.5$ $p=0.5$ |
| Genotype | $F_{1,48}=7.8$ $p=0.008$ | $F_{1,47}=0.1$ $p=0.7$ | $F_{1,48}=3.2$ $p=0.08$ | $F_{1,48}=3.4$ $p=0.07$ |
| Injury | $F_{1,48}=86.5$ $p<0.0001$ | $F_{1,47}=0.1$ $p=0.7$ | $F_{1,48}=92.9$ $p<0.0001$ | $F_{1,48}=18.4$ $p<0.0001$ |
| Time*genotype | $F_{1,48}=4.5$ $p=0.03$ | $F_{1,47}=5.7$ $p=0.06$ | $F_{1,48}=6.2$ $p=0.02$ | $F_{1,48}=0.002$ $p=1.0$ |
| Time*injury | $F_{1,48}=4.8$ $p=0.03$ | $F_{1,47}=0.7$ $p=0.4$ | $F_{1,48}=0.4$ $p=0.5$ | $F_{1,48}=0.3$ $p=0.6$ |
| Genotype*Injury | $F_{1,48}=2.2$ $p=0.1$ | $F_{1,47}=1.6$ $p=0.2$ | $F_{1,48}=0.9$ $p=0.4$ | $F_{1,48}=1.0$ $p=0.3$ |
| Time*genotype*injury | $F_{1,48}=1.9$ $p=0.2$ | $F_{1,47}=1.2$ $p=0.2$ | $F_{1,48}=0.2$ $p=0.7$ | $F_{1,48}=4.8$ $p=0.03$ |

**Table 4.** Three-way ANOVA assessing axonal bulb density.

|  | Ipsi | Contra |
| --- | --- | --- |
| DPI | $F_{1,65}=13.9$ $p=0.0004$ | $F_{1,65}=12.0$ $p=0.0009$ |
| Genotype | $F_{1,65}=0.7$ $p=0.4$ | $F_{1,65}=0.3$ $p=0.6$ |
| Injury | $F_{1,65}=8.5$ $p=0.005$ | $F_{1,65}=3.0$ $p=0.09$ |
| DPI*genotype | $F_{1,65}=0.7$ $p=0.4$ | $F_{1,65}=0.3$ $p=0.6$ |
| Time*injury | $F_{1,65}=8.4$ $p=0.005$ | $F_{1,65}=3.0$ $p=0.09$ |
| Genotype*Injury | $F_{1,65}=1.5$ $p=0.2$ | $F_{1,65}=0.1$ $p=0.7$ |
| Time*genotype*injury | $F_{1,65}=1.5$ $p=0.2$ | $F_{1,65}=0.1$ $p=0.7$ |

CHI produces a diffuse white matter injury that damages the axons, axonal transport, and myelin [21]. White matter damage is compared at 14 and 90 DPI between WT and MAPTKI mice. Axotomy in the corpus callosum produces axonal bulbs that are visualized using Bielschowsky’s silver stain (Fig. 3A) [21] [22]. At 14 or 90 DPI, axonal bulb number was similar in males and females, so they are combined (14DPI WT, F_1,13_=0.04 p=0.8, MAPTKI, F_1,22_=0.07 p=0.8; 90DPI, WT, F_1,12_= 0.001 p=1.0, MAPTKI, F_1,9_=0.001 p=1.0). Ipsilesional axonal bulb number has significant effects of injury and time*injury (Table 3). At 14 DPI, ipsilesional axonal bulb number significantly increased in injured WT mice that is no longer present at 90 DPI (Fig. 3B). Axonal bulbs did not significantly increase in the contralesional corpus callosum.

**Figure 3.**
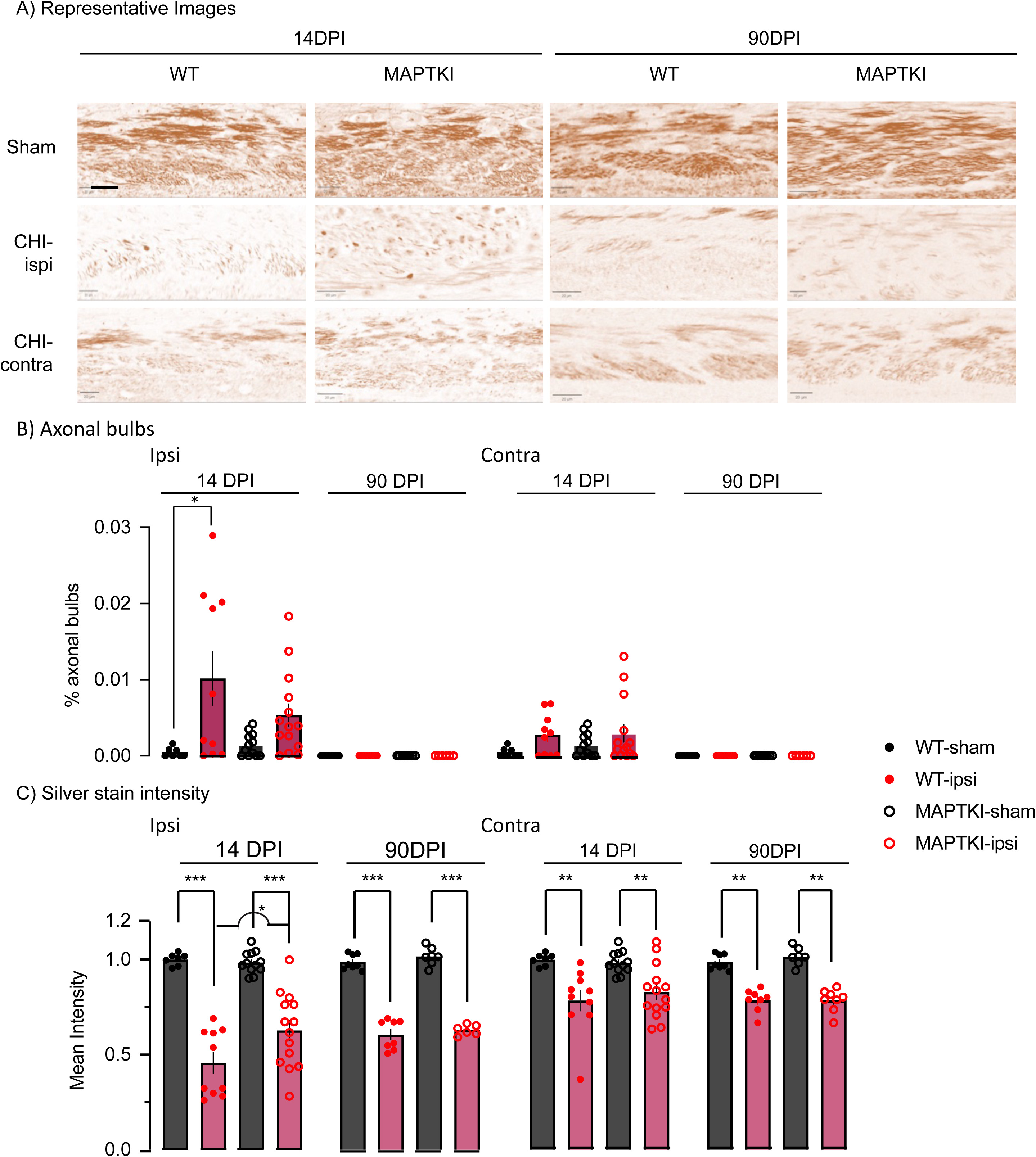
needs work

Bielschowsky’s silver stain also assesses overall axonal loss [59, 60]. At 14 DPI, stain intensity in MAPTKI corpus callosum has a significant sex effect yet no biologically significant pairwise comparisons. No other significant sex effects are noted (WT, 14DPI WT, F_1,13_=0.04 p=0.8, MAPTKI, F_1,22_=4.1 p=0.06; 90 DPI WT, F_1,12_= 0.2 p=0.7, MAPTKI, F_1,9_=0.001 p=1.0). The ipsilesional and contralesional corpus callosum has significant injury effects (Table 5). At 14 and 90 DPI, injury lowers staining intensity in the corpus callosum bilaterally. The injured ipsilesional WT corpus callosum has a lower staining intensity than MAPTKI at 14 DPI. The two strains have a similar loss of staining intensity at 90 DPI. These data suggest that the injured MAPTKI corpus callosum has less axonal loss at 14 DPI.

**Table 5.**
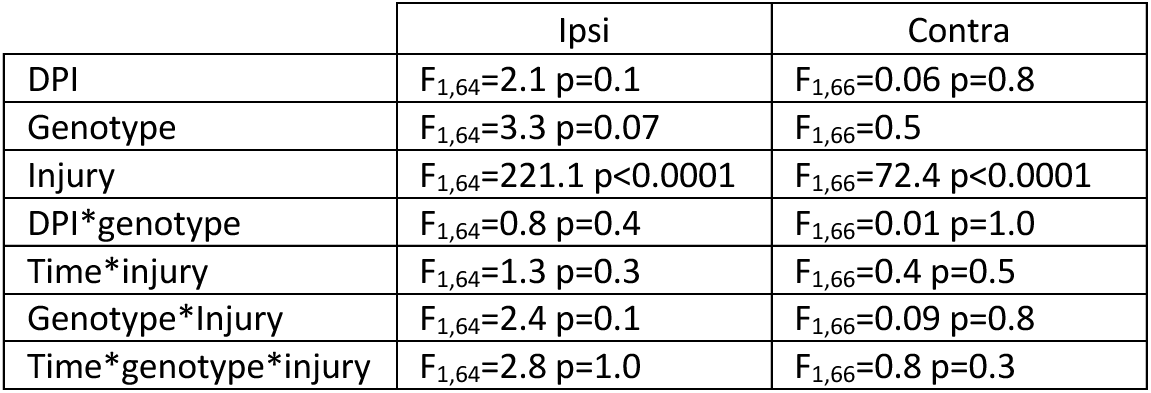
Three-Way ANOVA assessing corpus callosum axons stained with Bielschowsky’s silver stain.

|  | Ipsi | Contra |
| --- | --- | --- |
| DPI | $F_{1,64}=2.1$ $p=0.1$ | $F_{1,66}=0.06$ $p=0.8$ |
| Genotype | $F_{1,64}=3.3$ $p=0.07$ | $F_{1,66}=0.5$ |
| Injury | $F_{1,64}=221.1$ $p<0.0001$ | $F_{1,66}=72.4$ $p<0.0001$ |
| DPI*genotype | $F_{1,64}=0.8$ $p=0.4$ | $F_{1,66}=0.01$ $p=1.0$ |
| Time*injury | $F_{1,64}=1.3$ $p=0.3$ | $F_{1,66}=0.4$ $p=0.5$ |
| Genotype*Injury | $F_{1,64}=2.4$ $p=0.1$ | $F_{1,66}=0.09$ $p=0.8$ |
| Time*genotype*injury | $F_{1,64}=2.8$ $p=1.0$ | $F_{1,66}=0.8$ $p=0.3$ |

SMI-34 staining detects axonal loss or dephosphorylation of heavy neurofilament protein after TBI [61]. An effect of SMI-34 staining is compared in the WT or MAPTKI corpus callosum (Fig. 4). SMI-34 intensity lacked a significant sex effect at 14 or 90 DPI (14D WT F_1,14_=0.9 p=0.4, MAPTKI, F_1,14_=1.4 p=0.3; 90D WT F_1,13_=2.5 p=0.1, MAPTKI, F_1,8_=0.03 p=0.9). SMI-34 staining intensity in the ipsilesional and contralesional corpus callosum has significant effects of time, injury, and time*injury (Table 6). At 14 DPI, SMI-34 staining is lower in the injured ipsilesional WT and MAPTKI corpus callosum that is no longer present at 90 DPI (Fig. 4B). These data suggest that injured corpus callosum of WT and MAPTKI mice have a similar loss of phosphorylated heavy-chain neurofilament.

**Figure 4.**
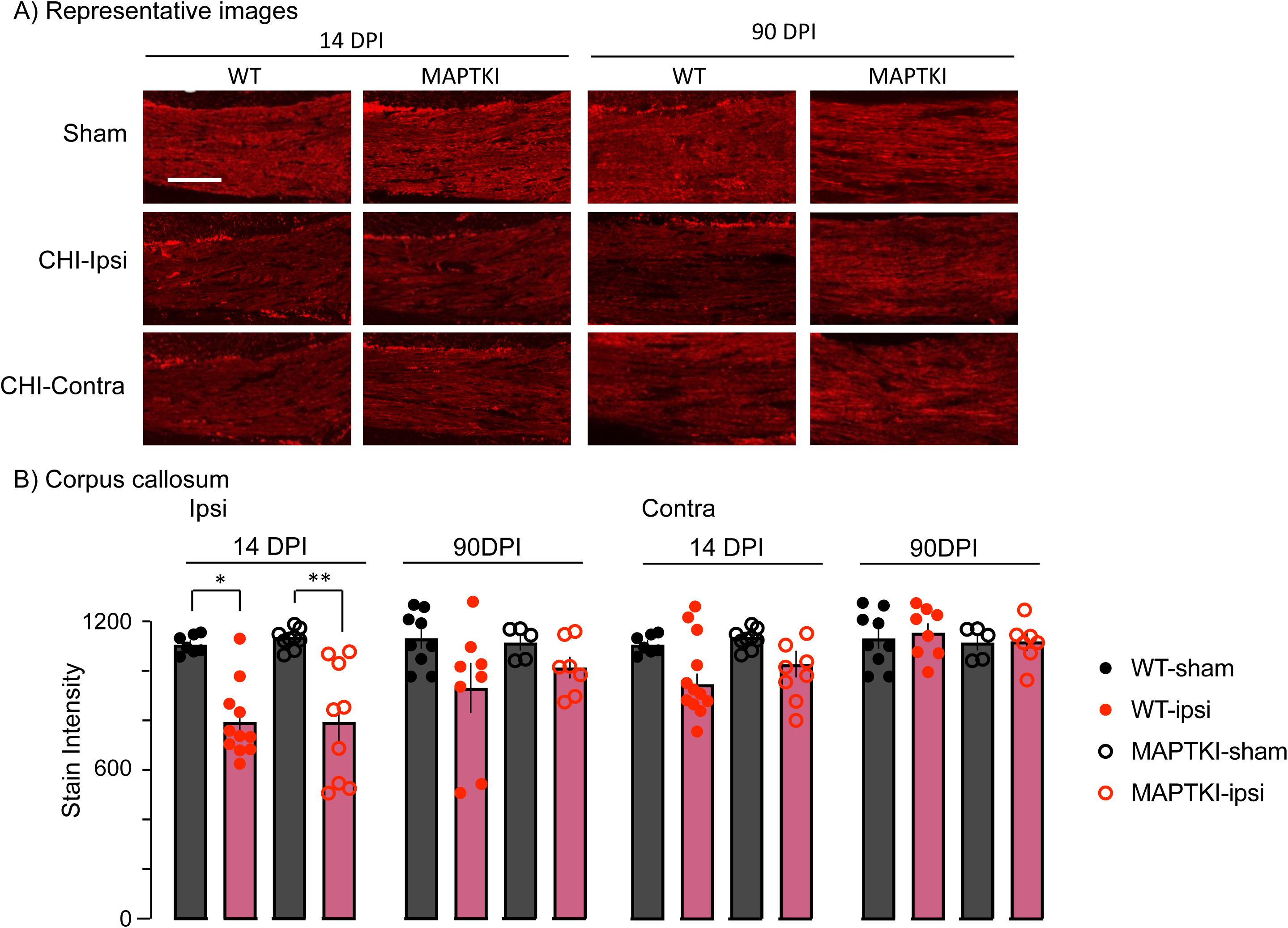
Similar loss of SMI-34 expression in the injured WT and MAPTKI corpus callosum. A) Representative images of SMI-34 expression in the WT and MAPTKI corpus callosum. Scale bar 50μm. B) Summary of SMI-34 expression in the WT and MAPTKI ipsilesional and contralesional corpus callosum. *p=0.003, **p=0.0004. n’s= 6-11.

**Table 6.**
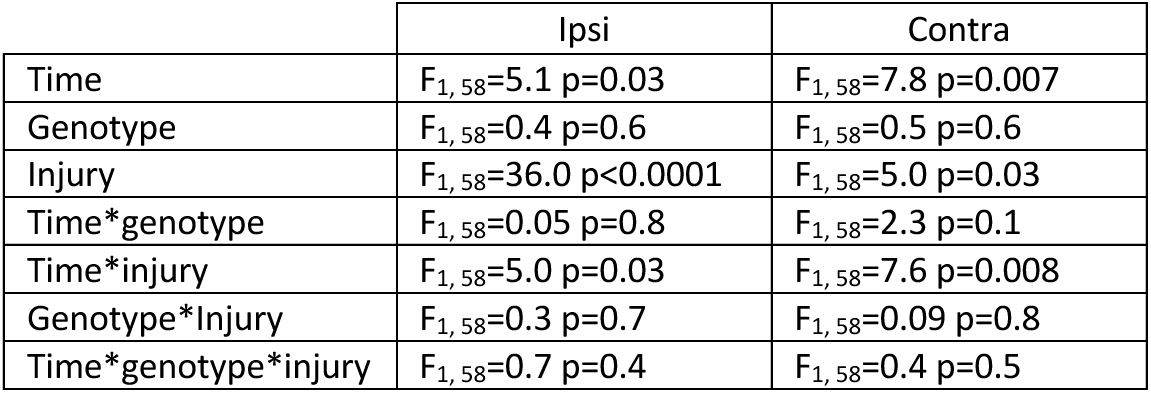
Three-Way ANOVA assessing corpus callosum axons stained with SMI-34.

|  | Ipsi | Contra |
| --- | --- | --- |
| Time | $F_{1,58}=5.1$ $p=0.03$ | $F_{1,58}=7.8$ $p=0.007$ |
| Genotype | $F_{1,58}=0.4$ $p=0.6$ | $F_{1,58}=0.5$ $p=0.6$ |
| Injury | $F_{1,58}=36.0$ $p<0.0001$ | $F_{1,58}=5.0$ $p=0.03$ |
| Time*genotype | $F_{1,58}=0.05$ $p=0.8$ | $F_{1,58}=2.3$ $p=0.1$ |
| Time*injury | $F_{1,58}=5.0$ $p=0.03$ | $F_{1,58}=7.6$ $p=0.008$ |
| Genotype*Injury | $F_{1,58}=0.3$ $p=0.7$ | $F_{1,58}=0.09$ $p=0.8$ |
| Time*genotype*injury | $F_{1,58}=0.7$ $p=0.4$ | $F_{1,58}=0.4$ $p=0.5$ |

TBI impairs fast axonal transport [62, 63]. Axons transport amyloid precursor protein (APP); impaired transport results in APP puncta [64]. APP puncta are measured in the corpus callosum of WT and MAPTKI mice at 14 and 90 DPI (Fig. 5). No sex effect was observed at either 14 or 90 DPI, (14 DPI, WT F1,15=0.7 p=0.4; MAPTKI, F1,13=0.008 p=0.9; 90 DPI, F1,15=0.06 p=0.8, F1,7=1.1 p=0.3). APP positive density has significant effects of time, genotype, injury, genotype*time, and time*Injury (Table 7). At 14 DPI, injury significantly increased APP puncta in the ipsilesional WT, but not MAPTKI, corpus callosum (Fig. 5B). At 90 DPI, APP puncta in injured corpus callosum returned to sham levels in both strains. These data suggest impaired axonal transport at 14 DPI in injured WT.

**Figure 5.**
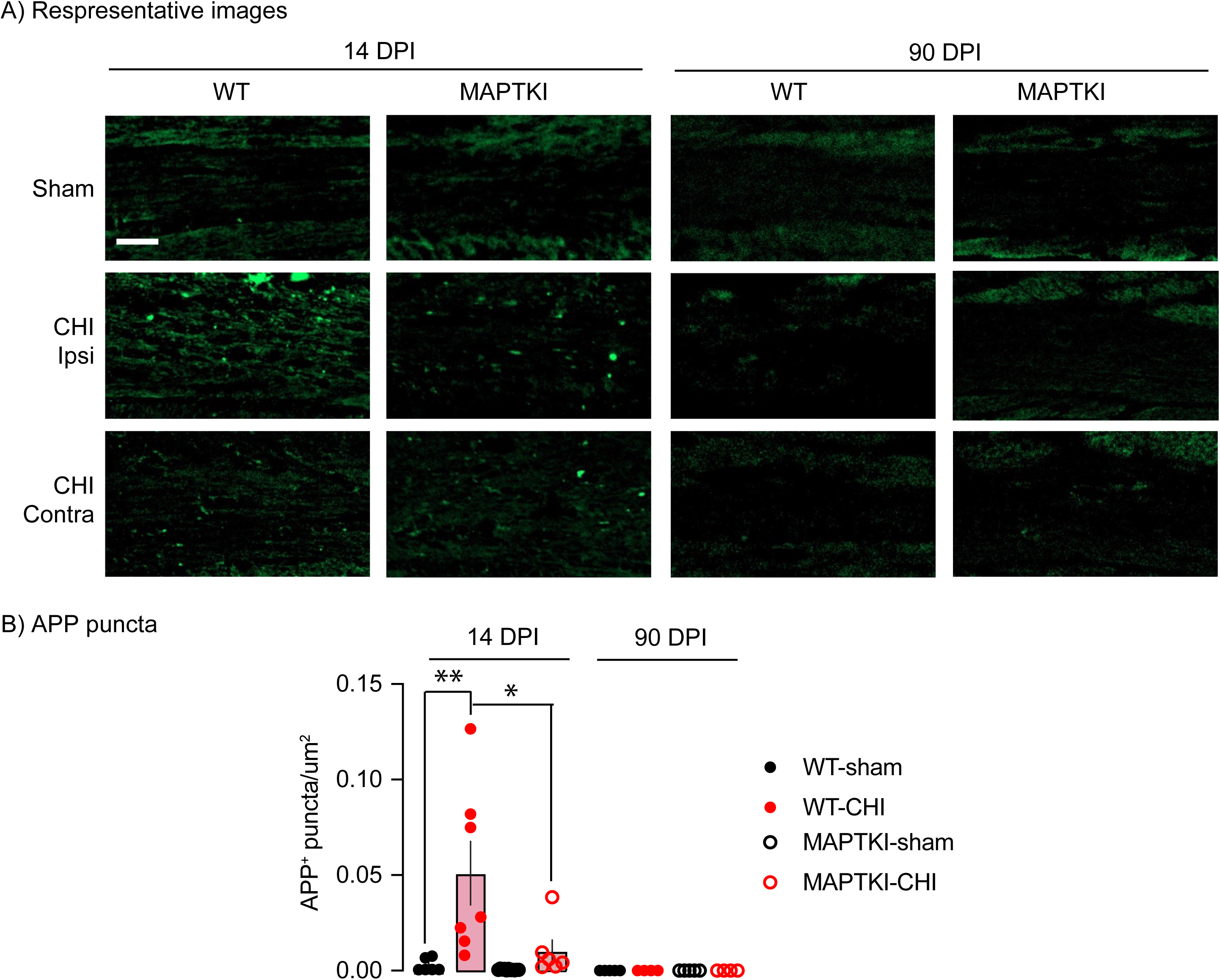
Accumulation of amyloid precursor protein (APP) in the injured WT corpus callosum. A) At 14 DP, APP puncta significantly increased in the corpus callosum of injured WT, but not MAPTKI mice. B) Summary of changes in APP puncta. **p<0.0001, *p=0.005.

**Table 7.** Three-way ANOVA analysis of corpus callosum APP puncta for effects of time, genotype or injury.

|  |  |
| --- | --- |
| Time | $F_{1,40}=9.5$ $p=0.004$ |
| Genotype | $F_{1,40}=6.7$ $p=0.01$ |
| Injury | $F_{1,40}=4.3$ $p=0.04$ |
| Genotype *time | $F_{1,38}=6.8$ $p=0.01$ |
| Time*Injury | $F_{1,40}=4.3$ $p=0.05$ |
| Genotype*Injury | $F_{1,40}=2.8$ $p=0.1$ |
| Time*Genotype*Injury | $F_{1,40}=2.8$ $p=0.1$ |

CHI produces myelin loss in WT at 14 DPI [23]. CHI WT or MAPTKI corpus callosum myelin content lacks a sex effect at 14 or 90 DPI (14 DPI, WT F_1,12_=0.05 p=0.8, MAPTKI F _1,14_ =1.4, p=0.3; 90 DPI, WT F _1,15_ =0.02 p=0.90, F _1,8_=0.2 p=0.7). Ipsilesional and contralesional myelin content has significant effects of injury, genotype*time and time*genotype*injury (Table 8). At 14 DPI, injury lowers myelin content bilaterally in the injured WT corpus callosum that returns to sham levels at 90 DPI (Fig. 6A,B). In contrast, myelin content decreased bilaterally in the MAPTKI corpus callosum at 90 DPI. These data suggest early myelin loss followed by remyelination in injured WT mice. In contrast, myelin loss is delayed in injured MAPTKI mice.

**Figure 6.**
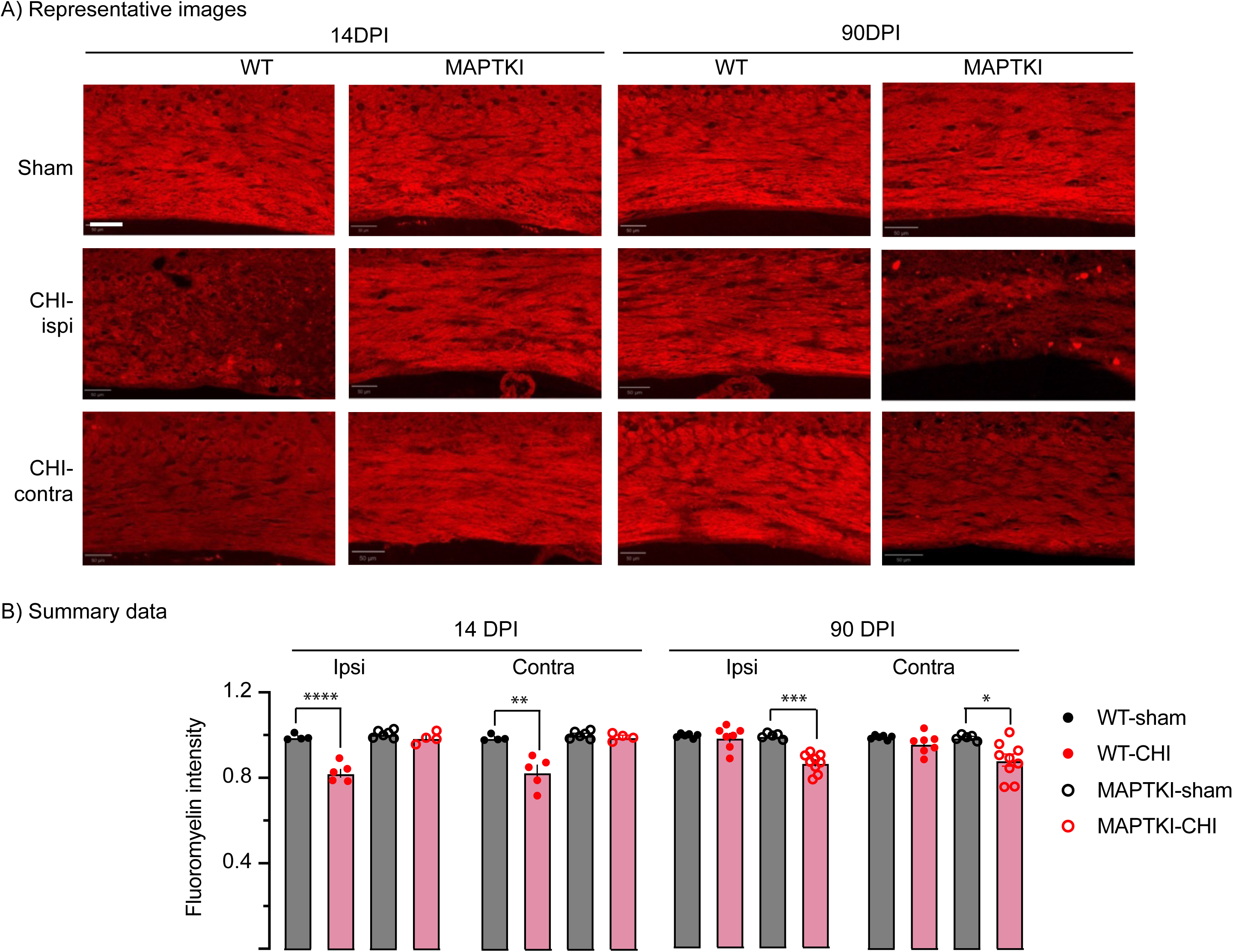
Myelin content in WT and MAPTKI mice. A) Representative images of fluoromyelin-stained ipsilesional and contralesional corpus callosum at 14 and 90 DPI. Scale bar 50μm. B) Summary of changes in fluoromyelin intensity at 14 or 90 DPI. *p=0.02, **p=0.008; ***p<0.0001. n’s=4-9.

**Table 8.** Three-way ANOVA analysis of myelin content.

|  | Ipsilesional | Contralesional |
| --- | --- | --- |
| Time | $F_{1,38}=2.1$ $p=0.15$ | $F_{1,38}=2.7$ $p=0.1$ |
| Genotype | $F_{1,38}=0.2$ $p=0.7$ | $F_{1,38}=0.001$ $p=0.9$ |
| Injury | $F_{1,38}=60.1$ $p<0.0001$ | $F_{1,38}=24.7$ $p<0.0001$ |
| Genotype *time | $F_{1,38}=51.4$ $p<0.0001$ | $F_{1,38}=17.3$ $p=0.0002$ |
| Time*Injury | $F_{1,38}=0.6$ $p=0.4$ | $F_{1,38}=1.4$ $p=0.2$ |
| Genotype*Injury | $F_{1,38}=1.5$ $p=0.2$ | $F_{1,38}=0.3$ $p=0.5$ |
| Time*Genotype*Injury | $F_{1,38}=41.0$ $p<0.0001$ | $F_{1,38}=13.3$ $p=0.0008$ |

Hyperphosphorylation promotes tau aggregation [65]. pTau levels are unchanged in uninjured WT assessed up to 24 months of age or in uninjured MAPTKI mice up to 24 months of age [21, 32]. CHI increased pTau expression in WT mice as assessed using AT8, PHF1 or S214 antibodies [21]. The same panel of antibodies assess changes in pTau^+^ cell density in the corpus callosum, cortex, and thalamus of injured WT and MAPTKI mice. AT8^+^ cell density has a significant sex effect yet produced no significant pairwise comparisons (Table 9). Therefore, results from males and females are combined. Corpus callosum has significant effects of injury, time*genotype and time*injury*genotype (Table 9). Injury increases AT8^+^ cell density in MAPTKI corpus callosum at 14 DPI that decreases at 90 DPI (Fig. 7B). At 90 DPI, AT8^+^ cell density increases in the injured WT corpus callosum. Cortex has a significant injury effect, yet with no significant pairwise effects (Fig. 7C). Thalamus has a significant injury effect AT8^+^ cell density increasing in WT mice at 14 DPI (Fig. 7D).

**Figure 7.**
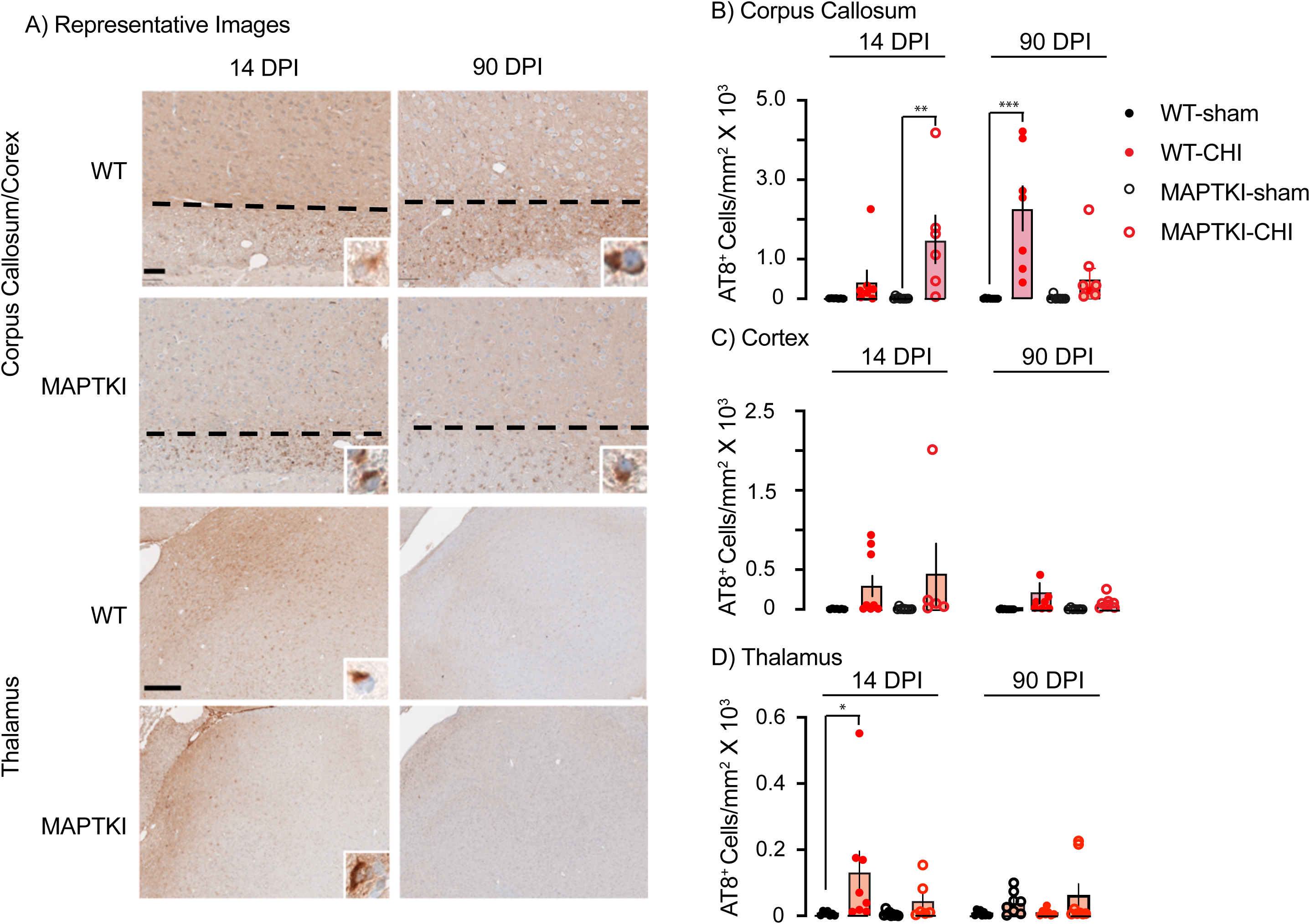
Injury increases pTau AT8 expression in corpus callosum and thalamus. A) Representative images of male or female AT8-stained cortex, corpus callosum and thalamus at 14 and 90 DPI. Dashed lines indicate the boundary of the cortex and corpus callosum. Insert shows higher magnification of AT8+ cells. Scale bars, corpus callosum, 50μm; thalamus, 250μm. B) Summary of changes in AT8^+^ cell density. *p=0.04, **p=0.01, ***p=0.0009. n’s= 4-7.

**Table 9.**
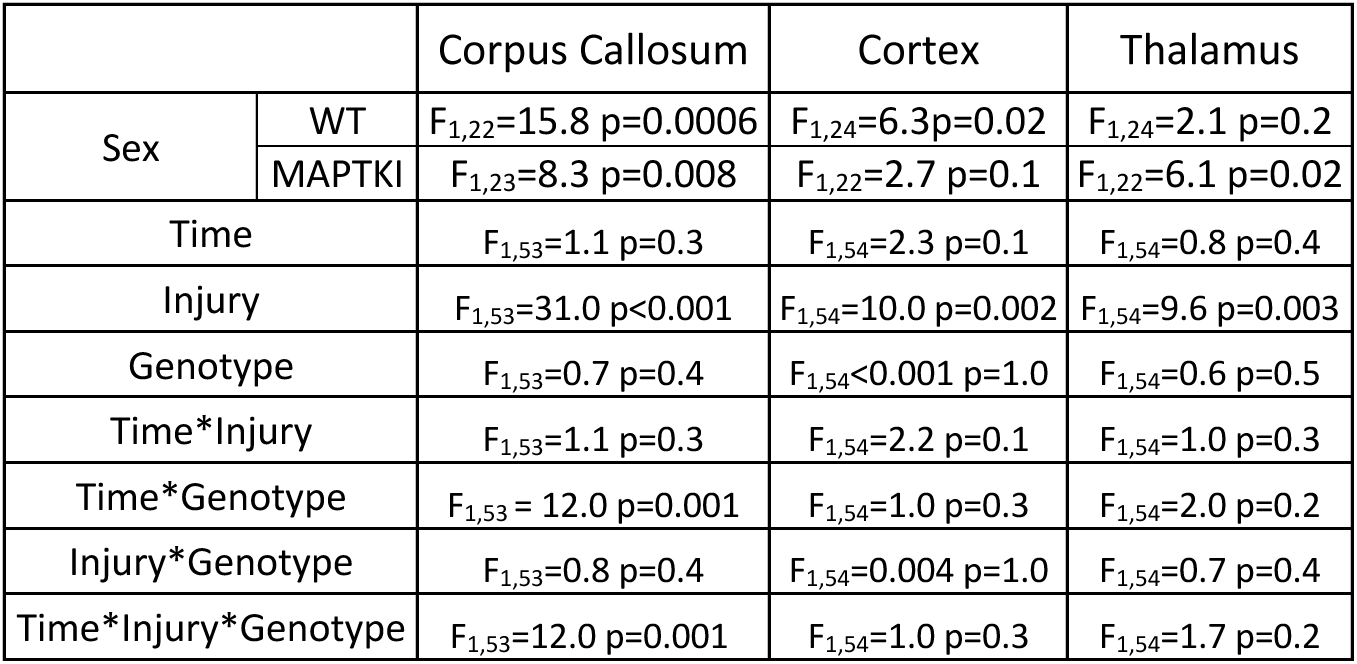
3-way-ANOVA analysis of AT8^+^ Cell Density.

PHF1^+^ cell density of males and females are analyzed separately since PHF1^+^ cell density has a significant sex effect in cortex, corpus callosum and thalamus (Table 10). Female cortical PHF1^+^ cell density has a significant injury effect with an increase of PHF1^+^ cell density at 14 DPI. PHF1^+^ cell density decreases, yet remained elevated, at 90 DPI (Fig. 8B). PHF1^+^ cell density is unchanged in the injured male cortex. In the thalamus, PHF1^+^ cell density in both sexes have significant effects of time (Table 10). Female thalamus has a significant genotype effect, while male thalamus has significant effects of injury and time*injury. At 14 DPI, PHF1^+^ cell density is elevated in the injured WT female thalamus which decreases to sham levels at 90 DPI. Male and female corpus callosum has significant effects of time and injury but lack biologically relevant pairwise comparisons.

**Figure 8.**
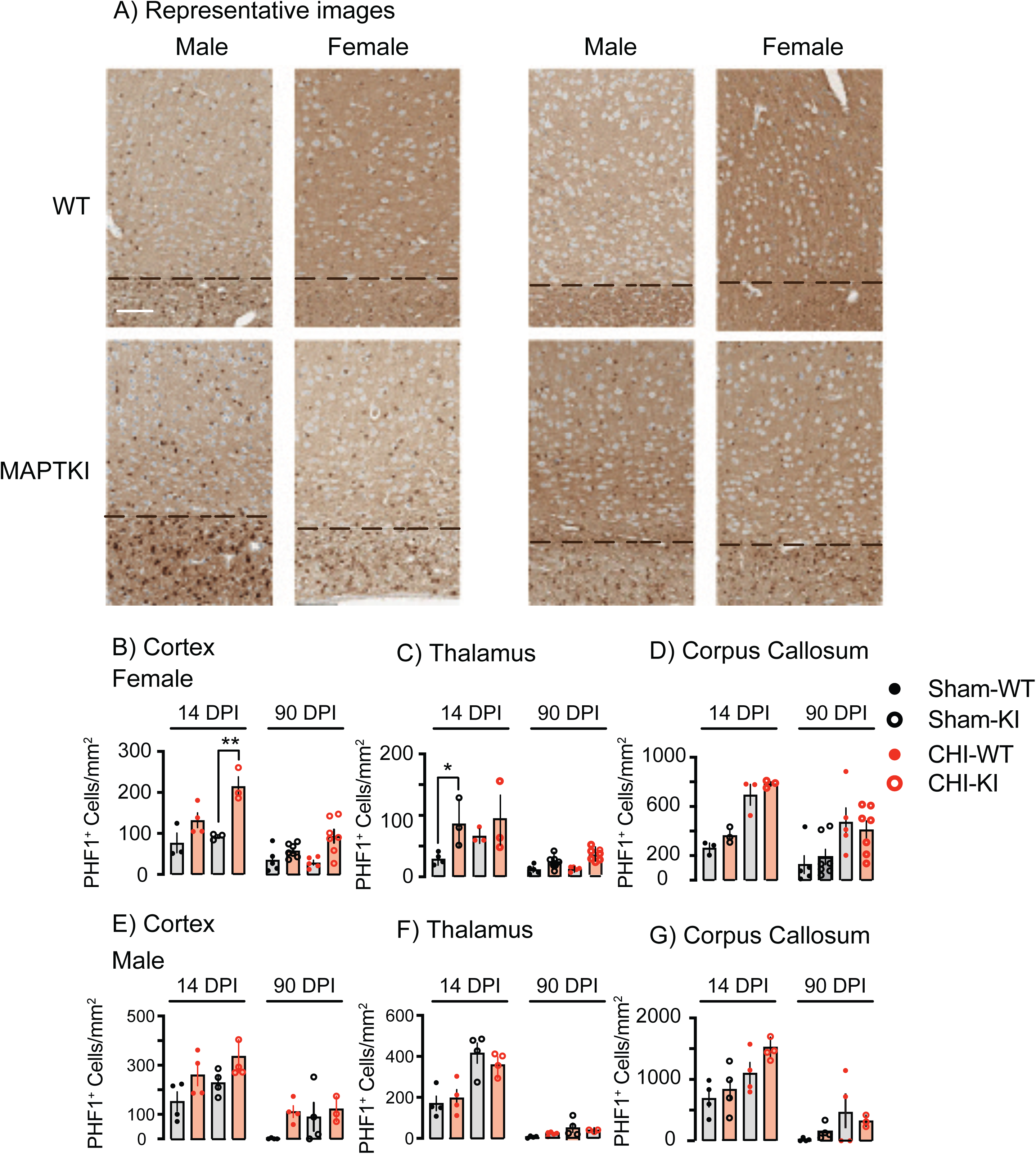
Injury increases PHF-1 pTau expression in the female corpus callosum. A) Representative images of male and female cortex, corpus callosum at 14 and 90DPI stained for PHF1 pTau. A dashed lines delineates the boundary of the corpus callosum and the cortex. Scale bar 50 μm. Summary of PHF1^+^ cell density in cortex (B,E). thalamus (C,F) and corpus callosum (D,G). *p=0.04, **p=0.004, ***p=0.003. n’s=3-5.

**Table 10.** 3-Way-ANOVA analysis of PHF1^+^ Cell Density.

|  | Genotype | Corpus Callosum | Thalamus | Cortex |
| --- | --- | --- | --- | --- |
| Sex | WT | $F_{1,24}=3.0$ $p=1.0$ | $F_{1,25}=70.5$ $p<0.0001$ | $F_{1,25}=6.5$ $p=0.02$ |
| | MAPTKI | $F_{1,27}=13.1$ $p=0.001$ | $F_{1,27}=46.0$ $p<0.0001$ | $F_{1,27}=19.5$ $p=0.0001$ |

Table 10. 3-Way-ANOVA analysis of PHF1<sup>+</sup> Cell Density
|  | Sex | Corpus Callosum | Thalamus | Cortex |
| --- | --- | --- | --- | --- |
| Time | Male | $F_{1,23}=48.4$ $p<0.0001$ | $F_{1,24}=151.1$ $p<0.0001$ | $F_{1,24}=26.37$ $p<0.0001$ |
| | Female | $F_{1,28}=13.7$ $p=0.0009$ | $F_{1,28}=36.8$ $p<0.0001$ | $F_{1,28}=49.6$ $p<0.0001$ |
| Injury | Male | $F_{1,23}=13.3$ $p=0.001$ | $F_{1,24}=34.8$ $p<0.0001$ | $F_{1,24}=3.7$ $p=0.07$ |
| | Female | $F_{1,28}=33.8$ $p<0.0001$ | $F_{1,28}=2.8$ $p=0.1$ | $F_{1,24}=23.3$ $p<0.0001$ |
| Genotype | Male | $F_{1,23}=1.2$ $p=0.3$ | $F_{1,24}=0.03$ $p=0.9$ | $F_{1,24}=8.7$ $p=0.007$ |
| | Female | $F_{1,28}=0.6$ $p=0.5$ | $F_{1,28}=13.6$ $p=0.001$ | $F_{1,24}=18.8$ $p=0.0002$ |
| Time*Injury | Male | $F_{1,23}=0.8$ $p=0.4$ | $F_{1,24}=18.5$ $p=0.0002$ | $F_{1,24}=3.7$ $p=0.8$ |
| | Female | $F_{1,28}=1.4$ $p=0.2$ | $F_{1,28}=1.4$ $p=0.3$ | $F_{1,24}=12.2$ $p=0.002$ |
| Time*Genotype | Male | $F_{1,23}=1.3$ $p=0.3$ | $F_{1,24}=0.008$ $p=0.9$ | $F_{1,24}=0.2$ $p=0.7$ |
| | Female | $F_{1,28}=0.6$ $p=0.4$ | $F_{1,28}=2.7$ $p=0.1$ | $F_{1,24}=0.07$ $p=0.8$ |
| Injury*Genotype | Male | $F_{1,23}=0.03$ $p=0.9$ | $F_{1,24}=1.6$ $p=0.2$ | $F_{1,24}=1.6$ $p=0.2$ |
| | Female | $F_{1,28}=0.3$ $p=0.6$ | $F_{1,28}=0.5$ $p=0.5$ | $F_{1,24}=6.5$ $p=0.02$ |
| Time*Injury*Genotype | Male | $F_{1,23}=1.1$ $p=0.3$ | $F_{1,24}=0.06$ $p=0.8$ | $F_{1,24}=0.2$ $p=0.7$ |
| | Female | $F_{1,28}=0.2$ $p=0.7$ | $F_{1,28}=1.2$ $p=0.3$ | $F_{1,24}=0.3$ $p=0.6$ |

S214^+^ cell density has significant sex effects in cortex, thalamus, and corpus callosum therefore, females and males are analyzed separately (Table 11). Female cortical S214^+^ cell density has significant effects of time, injury, and time*injury (Table 11). At 14 DPI, S214^+^ cell density increases female WT and MAPTKI cortex that is no longer present at 90 DPI (Fig. 9B,E). Male thalamic S214^+^ cell density has significant effects of injury and time*genotype yet with no biologically significant pairwise effects. Female S214^+^ cell density in corpus callosum had significant effects of time, injury, time*injury (Table 11). S214^+^ cell density in the injured female WT corpus callosum increases at 14 DPI that decreases at 90 DPI (Fig. 9D). Male cortical S214^+^ cell density had significant effects of time, injury, and time*injury (Table 11). Injured WT and MAPTKI mice increased S214^+^ cell density at 14 DPI (Fig. 9E). Male thalamic S214^+^ cell density had a significant injury effect with no biological relevant pairwise comparisons. Male S214^+^ cell density in corpus callosum has a significant injury effect time*genotype. At 14 DPI, injury increases male S214^+^ cell density in corpus callosum that decreases at 90 DPI (Fig. 9G). Thioflavin-S (thio-S) stains β -sheet structures in toxic protein aggregates (ref). Injured WT mice at 180 DPI increase thalamic thio-S^+^ cell density (Havlicek). Thio-S^+^ cell density in a mixed cohort of female WT and MAPTKI mice is assessed in the corpus callosum, thalamus, and cortex of male and at 90 DPI. Thio-S^+^ cell density in the corpus callosum has a significant effect of genotype, injury and genotype*injury (Table 13). Injury increases thio-S^+^ cell density in the corpus callosum of MAPTKI but not WT, mice (Fig. 10B). Both cortical and thalamic thio-S cell density have a significant effect of injury, but not genotype or genotype*injury (Table 12). Injury increases cortical thio-S cell density in both WT and MAPTKI mice (Fig. 10B). Only WT mice increase thalamic thio S^+^ cell density (Fig. 10B).

**Figure 9.**
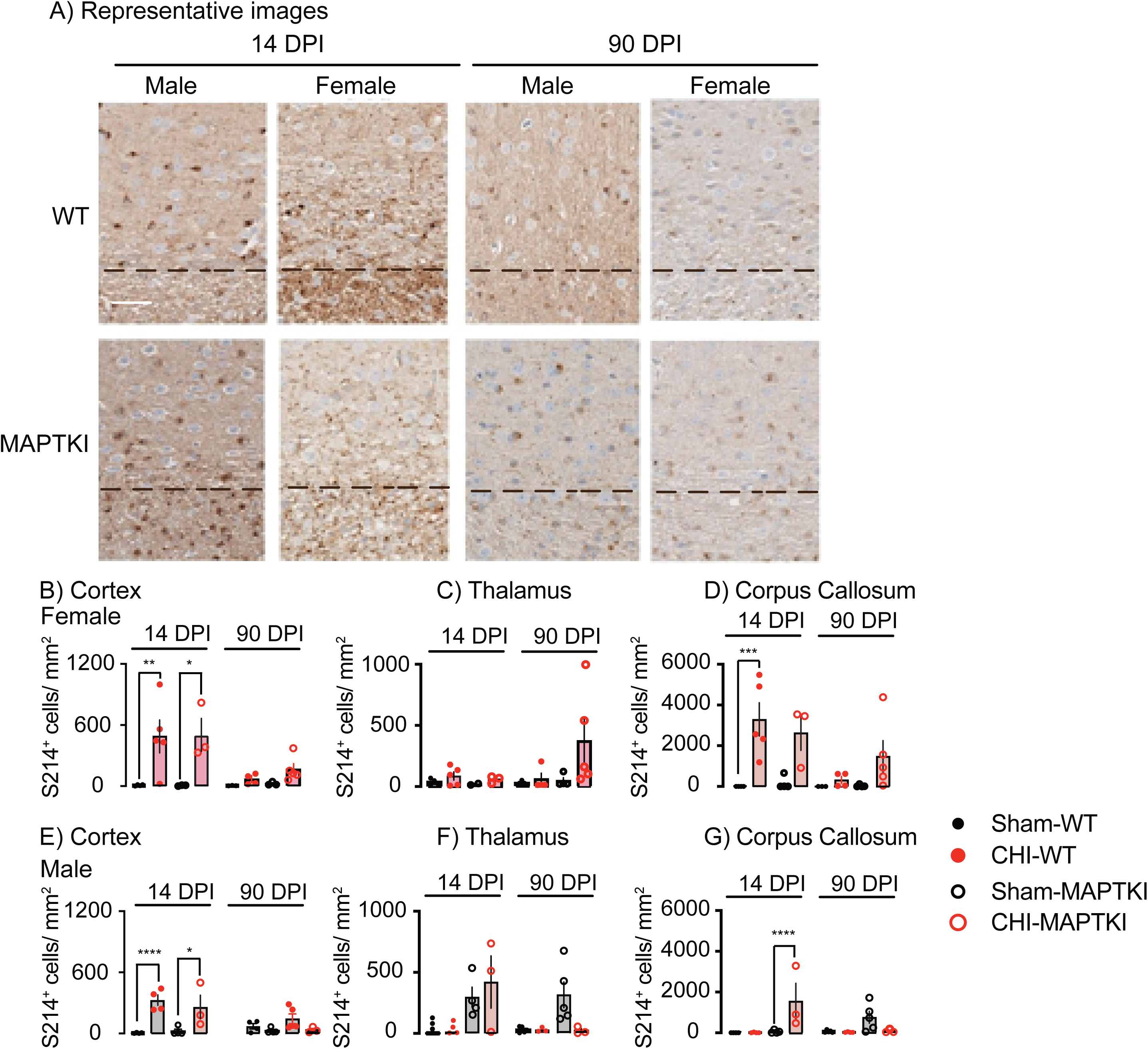
Injury increases S214^+^ cell density at 14 DPI. A) Representative images of male and female cortex, corpus callosum at 14 and 90DPI stained for pTau S214. Scale bar 50 μm. S214^+^ cell density is assessed in female and male cortex (A,D), thalamus (B,D) and corpus callosum (C,E). At 14 DPI, injury increases S214^+^ cell density in WT male and female cortex and corpus callosum. Injury also increases S214^+^ cell density in MAPTKI corpus callosum *p=0.04, **p=0.007, ***p=0.003, ****p<0.0005. n’s=3-5

**Figure 10.**
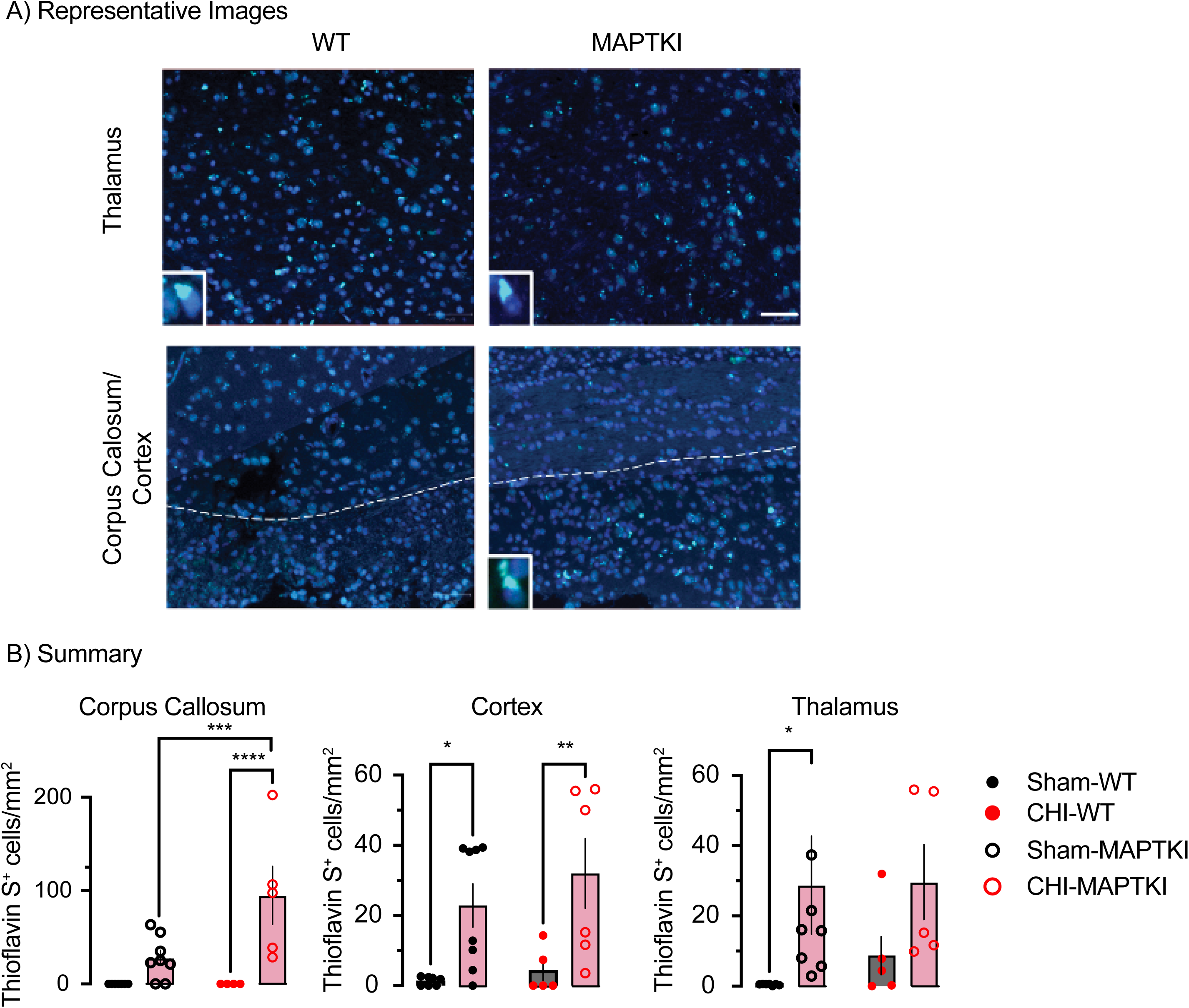
Injury increases thio-S^+^ cell density. **A**) Representative images of thalamus, cortex, and corpus callosum stained with Thio S (green) and DAPI (blue). Higher power insets show characteristic perinuclear staining of Thio S. Dotted lines delineate the boundary between corpus callosum and cortex. Scale bar, 50 μm. B) Summary of changes in Thio-S^+^ cell density. *p<0.05, **p=0.007, ***p=0.002, ****p=0.0005. n’s=4-8.

**Table 11.** 3-Way-ANOVA analysis of S214^+^ Cell Density.

|  | Cortex |  | Corpus Callosum |  | Thalamus |  |
| --- | --- | --- | --- | --- | --- | --- |
| DPI | 14 | 90 | 14 | 90 | 14 | 90 |
| Sex | $F_{1,22}=2.0$<br>$p=0.17$ | $F_{1,25}=0.0001$<br>$p=1.0$ | $F_{1,22}=8.3$<br>$p=0.009$ | $F_{1,25}=0.9$<br>$p=0.4$ | $F_{1,22}=6.5$<br>$p=0.02$ | $F_{1,25}=0.13$<br>$p=0.7$ |

Table 11 3-Way-ANOVA analysis of S214<sup>+</sup> Cell Density
|  | Sex | Cortex | Corpus Callosum | Thalamus |
| --- | --- | --- | --- | --- |
| Time | Male | $F_{1,23}=7.3$ $p=0.01$ | $F_{1,24}=1.0$ $p=0.3$ | $F_{1,23}=2.0$ $p=0.2$ |
| | Female | $F_{1,22}=7.7$ $p=0.01$ | $F_{1,23}=6.0$ $p=0.02$ | $F_{1,22}=1.2$ $p=0.3$ |
| Injury | Male | $F_{1,22}=23.4$ $p<0.0001$ | $F_{1,24}=10.0$ $p=0.004$ | $F_{1,23}=18.0$ $p=0.0003$ |
| | Female | $F_{1,22}=18.9$ $p=0.0003$ | $F_{1,23}=21.3$ $p=0.0001$ | $F_{1,22}=2.7$ $p=0.11$ |
| Genotype | Male | $F_{1,23}=2.4$ $p=0.1$ | $F_{1,24}=1.1$ $p=0.3$ | $F_{1,23}=0.5$ $p=0.5$ |
| | Female | $F_{1,22}=0.2$ $p=0.7$ | $F_{1,23}=0.1$ $p=0.8$ | $F_{1,22}=1.1$ $p=0.3$ |
| Time*Injury | Male | $F_{1,23}=13.0$ $p=0.002$ | $F_{1,24}=1.3$ $p=0.3$ | $F_{1,23}=3.4$ $p=0.07$ |
| | Female | $F_{1,22}=8.1$ $p=0.009$ | $F_{1,23}=6.2$ $p=0.02$ | $F_{1,22}=1.0$ $p=0.3$ |
| Time* Genotype | Male | $F_{1,23}=0.6$ $p=0.4$ | $F_{1,24}=9.0$ $p=0.006$ | $F_{1,23}=3.3$ $p=0.08$ |
| | Female | $F_{1,22}=0.2$ $p=0.7$ | $F_{1,23}=1.2$ $p=0.3$ | $F_{1,22}=2.0$ $p=0.2$ |
| Injury* Genotype | Male | $F_{1,23}=1.3$ $p=0.3$ | $F_{1,24}=1.3$ $p=0.3$ | $F_{1,23}=0.6$ $p=0.5$ |
| | Female | $F_{1,22}=0.09$ $p=0.8$ | $F_{1,23}=0.07$ $p=0.8$ | $F_{1,22}=1.1$ $p=0.3$ |
| Time* Injury* Genotype | Male | $F_{1,23}=0.02$ $p=0.7$ | $F_{1,24}=8.2$ $p=0.008$ | $F_{1,23}=3.1$ $p=0.09$ |
| | Female | $F_{1,22}=0.08$ $p=0.8$ | $F_{1,23}=1.1$ $p=0.3$ | $F_{1,22}=1.2$ $p=0.3$ |

**Table 12.** Two-way ANOVA analysis of Thio S^+^ cell density.

|  | Corpus Callosum | Thalamus | Cortex |
| --- | --- | --- | --- |
| Genotype | $F_{1,20}=5.4$ $p=0.03$ | $F_{1,21}=0.19$ $p=0.7$ | $F_{1,22}=1.0$ $p=0.3$ |
| Injury | $F_{1,20}=18.2$ $p=0.0004$ | $F_{1,21}=5.4$ $p=0.02$ | $F_{1,22}=15.9$ $p=0.0006$ |
| Genotype*Injury | $F_{1,20}=5.4$ $p=0.03$ | $F_{1,21}=0.12$ $p=0.7$ | $F_{1,22}=0.2$ $p=0.6$ |

**Table 13.** Distance traveled (meters) on Barnes maze.

| DPI | 14 |  | 90 |  |
| --- | --- | --- | --- | --- |
| Training day | 1 | 4 | 1 | 4 |
| WT-Sham | 2.3±0.3 | 0.8±0.2 | 3.8±0.4 | 1.7±0.4 |
| WT-CHI | 4.1±0.5 | 2.4±0.5* | 4.7±0.7 | 1.2±0.1 |
| MAPTKI-sham | 2.8±0.4 | 0.8±0.1 | 3.1±2.3 | 1.0±0.1 |
| MAPTKI-CHI | 2.2±0.3 | 0.9±0.1 | 3.5±0.4 | 1.2±0.1 |

WT and MAPTKI mice are tested at 14 and 90 DPI to test if injury produces differing behavioral deficits. Injured WT mice at 14 DPI are impaired in acquiring and retaining the location of a Barnes maze escape hole [20, 23]. It is unknown if these deficits persist at later times or are present in MAPTKI mice. Therefore, sham or injured WT and MAPTKI mice are tested on Barnes maze at 14 and 90 DPI. At 14 DPI, distance traveled has significant effects of day, genotype, injury, and genotype*injury (Tables 13, 14). On training day 1 at 14 DPI, injured WT mice traveled more than injured MAPTKI mice (Table 14). At 90 DPI, distance traveled has significant effects of day and genotype, yet with no significant pairwise comparisons. Sex affects Barnes maze performance [66]. At 14 DPI, WT latency to find the escape hole has a significant sex effect (WT, F_1,15_=11.4 p=0.004; MAPTKI, F_1, 14_= 3.2 p=0.09). On training day 1, the latency of WT sham females is shorter than males (p=0.03). This sex effect is no longer evident on trials 2 through 4. MAPTKI mice lacked any sex effect on any trials. WT or MAPTKI males and females are analyzed together. Barnes maze latency has a significant effect of trial, genotype, injury and trial*genotype*injury (Table 15). Both WT and MAPTKI mice lowered time to find the escape box between trials 1 and 4 (Fig. 11A). Latency of injured WT or MAPTKI mice on the 4^th^ and final training is greater than shams (Fig. 11A). This suggests a similarly mild impairment on Barnes maze for WT and MAPTKI mice. The following day, a probe trial tests retention of the escape hole location. Probe trial results from males and females are combined since WT mice lack a sex effect and the significant sex effect of MAPTKI mice lacks biologically relevant pairwise comparisons (WT, F_1,19_= 0.6 p=0.6; MAPTKI, F_1,15_= 4.5, p=0.05). Entrances in the probe trial has significant effects of injury and genotype*injury (Table 16). Injury impairs WT, but not MAPTKI mice, on probe trial (Fig. 11C). Sham and injured MAPTKI mice spend a similar time in target quadrant. These data suggest that, at 14 DPI, injured MAPTKI mice retain the escape box location while injured WT mice are impaired.

**Figure 11.**
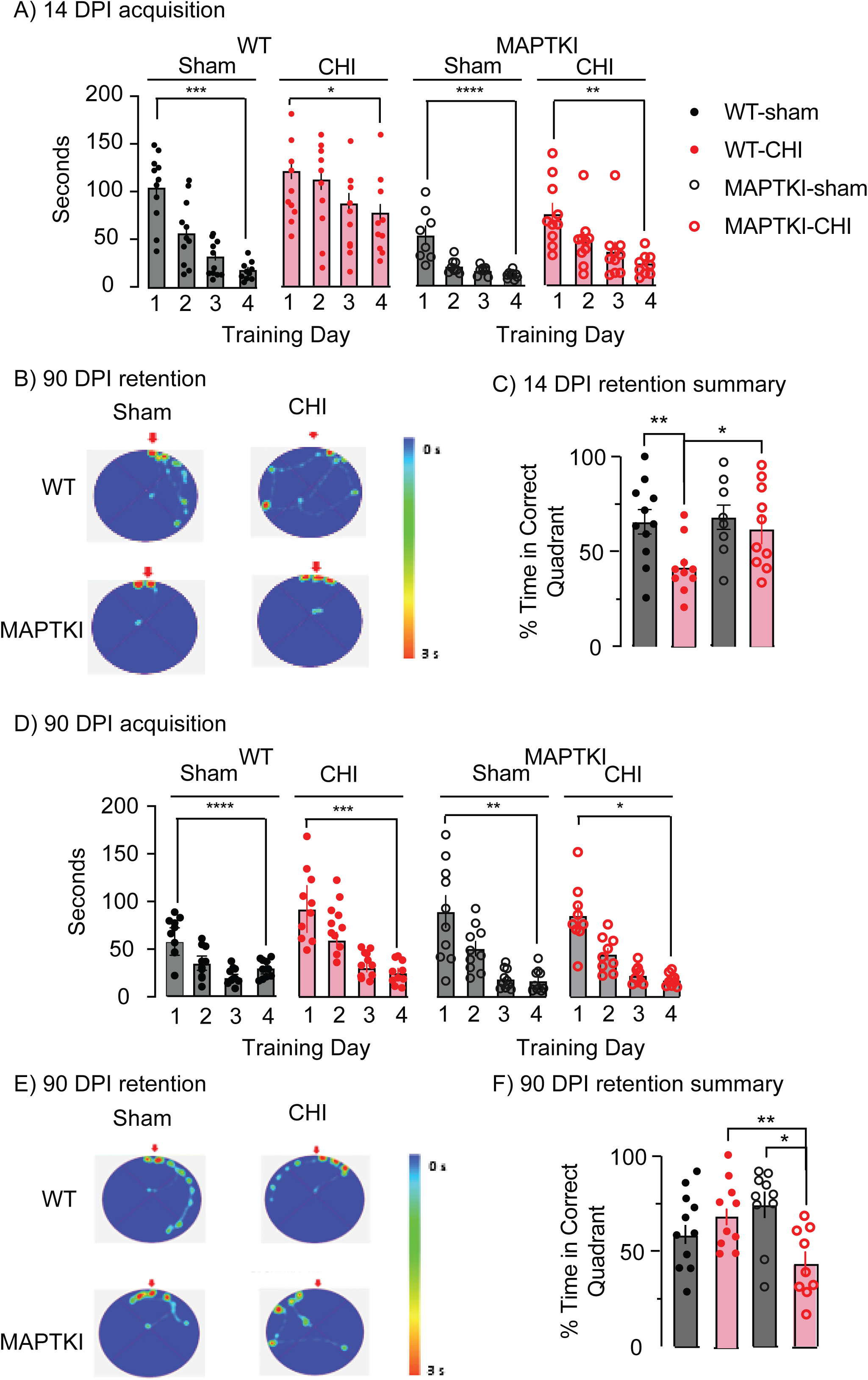
WT and MAPTKI mice differ in Barnes acquisition and retention. A) Summary of Barnes maze acquisition at 14 DPI. Training curves of WT (left) and MAPTKI (right) mice are graphed separately. All groups lowered latency between training 1 to 4. ****p=0.006, ***p=0.02, **p=0.05, *p=0.04. (B) Heat map of mouse location on probe trial on training day 5. An arrow indicates the escape hole location. (C) Summary of Barnes maze retention at 14 DPI. * p<0.05, **p<0.01. D) Summary of Barnes maze acquisition at 90 DPI. All groups lowered latency between training days 1 to 4. ****p=0.006, ***p=0.05, **p=0.03, *p=0.02. E) Heat map of mouse location on probe trial on training day 5. An arrow indicates the escape hole location. (F) Summary of Barnes maze retention at 90 DPI. WT mice retained the escape hole location while MAPTKI did not. **p=0.001, *p=0.01. n’s=9-11.

**Table 14.** Three-way ANOVA analysis of distance traveled on Barnes maze.

|  | 14 DPI | 90 DPI |
| --- | --- | --- |
| Training Day | $F_{1,211}=50.3$ $p<0.0001$ | $F_{1,301}=68.2$ $p<0.0001$ |
| Genotype | $F_{1,211}=9.8$ $p=0.002$ | $F_{1,301}=5.0$ $p=0.03$ |
| Injury | $F_{1,211}=9.1$ $p=0.003$ | $F_{1,301}=0.6$ $p=0.5$ |
| Genotype *day | $F_{1,211}=0.04$ $p=0.8$ | $F_{1,301}=0.9$ $p=0.3$ |
| Day*Injury | $F_{1,211}=0.4$ $p=0.5$ | $F_{1,301}=1.2$ $p=0.3$ |
| Genotype*Injury | $F_{1,211}=17.0$ $p<0.0001$ | $F_{1,301}=0.03$ $p=0.9$ |
| Day*Genotype*Injury | $F_{1,211}=4.0$ $p=0.4$ | $F_{1,301}=1.0$ $p=0.3$ |

**Table 15.** 3-way ANOVA analysis of Barnes Maze Training.

| DPI | 14 | 90 |
| --- | --- | --- |
| Trial | $F_{2.67, 133.4}=36.3$ $p<0.001$ | $F_{1.6, 57.7}=82.4$ $p<0.0001$ |
| Genotype | $F_{1, 50}=36.2$ $p<0.001$ | $F_{1,36} =0.1$ $p=0.7$ |
| Injury | $F_{1, 50}= 23.8$ $p<0.001$ | $F_{1,36} = 0.1$ $p=0.7$ |
| Trial*Genotype | $F_{3, 150}=2,2$ $p=0.09$ | $F_{3,108}=1.3$ $p=0.3$ |
| Trial*Injury | $F_{3, 150}=2,2$ $p=0.13$ | $F_{3,108}=1.4$ $p=0.2$ |
| Genotype*Injury | $F_{1, 50}=3.6$ $p=0.07$ | $F_{1,36}=1.1$ $p=0.3$ |
| Trial*Genotype*Injury | $F_{3,150}=2.7$ $p=0.05$ | $F_{3,108}=0.6$ $p=0.6$ |

**Table 16.**
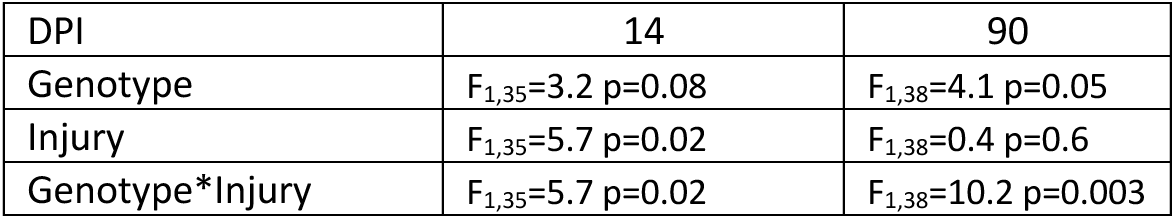
Two Way ANOVA analysis of Barnes Maze Probe Trial.

At 90 DPI, males and females are combined since MAPTKI or WT latency lacks a sex effect (WT, F_1,19_= 0.5 p=0.5; MAPTKI, F_1,15_= 0.02, p=0.9). At 90 DPI, Barnes maze latency has a trial significant effect (Table 15). At 90DPI, all four groups lower latency between days 1 and 4 (Fig. 11D). Males and females are combined on the Barnes probe trial at 90 DPI since WT mice lack a sex effect while MAPTKI mice have a sex affect with no biologically significant pairwise effects (WT, F_1,19_=3.0 p=0.09; MAPTKI, F_1,15_= 6.0, p=0.03). At 90 DPI, probe trial has significant effects of injury and genotype*injury (Table 16). Sham and injured WT mice spend similar time in the target quadrant (Fig. 11E, F). In contrast, injured MAPTKI mice are the target quadrant for significantly less time than shams (Fig. 11E, F). These data suggest that, at 90DPI, injured MAPTKI mice have impaired retention of Barnes maze. Memory impairments in injured WT mice at 14 DPI resolve by 90 DPI. In contrast, injured MAPTKI mice develop a memory deficit between 14 and 90 DPI.

Mice are also assessed on APA, a task having higher cognitive demand that Barnes maze [49]. Distance traveled and speed assesses motor performance in habituation, acquisition, and probe trials (Table 17). Results from APA acquisition trials at 14 or 90 DPI from male and female mice are combined since sex has no affect APA entrances for WT or MAPTKI mice (14 WT, F_1,23_=0.9 p=0.4, MAPTKI, F_1,15_=1.1 p=0.3; 90, WT, F_1,31_=11.5 p=0.69; MAPTKI, F_1,14_=1.9 p=0.2). During habituation, WT and MAPTKI mice have similar motor performance (Tables 17,18).

**Table 17.**
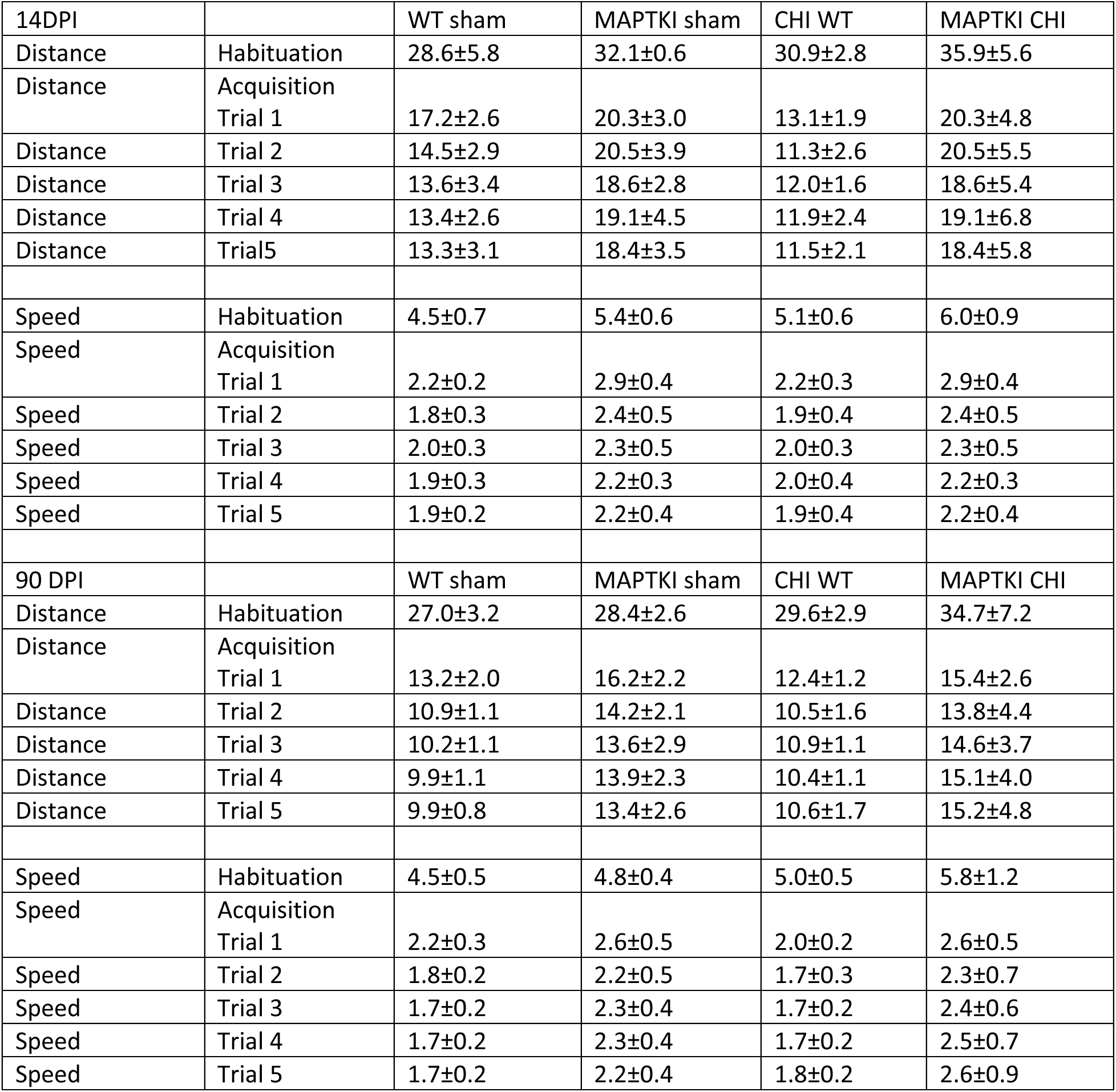
Distance (meters) and speed (cm/s) on active place avoidance habituation and acquisition.

|  |  |  |  |  |  |
| --- | --- | --- | --- | --- | --- |
| 14DPI |  | WT sham | MAPTKI sham | CHI WT | MAPTKI CHI |
| Distance | Habituation | 28.6±5.8 | 32.1±0.6 | 30.9±2.8 | 35.9±5.6 |
| Distance | Acquisition Trial 1 | 17.2±2.6 | 20.3±3.0 | 13.1±1.9 | 20.3±4.8 |
| Distance | Trial 2 | 14.5±2.9 | 20.5±3.9 | 11.3±2.6 | 20.5±5.5 |
| Distance | Trial 3 | 13.6±3.4 | 18.6±2.8 | 12.0±1.6 | 18.6±5.4 |
| Distance | Trial 4 | 13.4±2.6 | 19.1±4.5 | 11.9±2.4 | 19.1±6.8 |
| Distance | Trial5 | 13.3±3.1 | 18.4±3.5 | 11.5±2.1 | 18.4±5.8 |
| Speed | Habituation | 4.5±0.7 | 5.4±0.6 | 5.1±0.6 | 6.0±0.9 |
| Speed | Acquisition Trial 1 | 2.2±0.2 | 2.9±0.4 | 2.2±0.3 | 2.9±0.4 |
| Speed | Trial 2 | 1.8±0.3 | 2.4±0.5 | 1.9±0.4 | 2.4±0.5 |
| Speed | Trial 3 | 2.0±0.3 | 2.3±0.5 | 2.0±0.3 | 2.3±0.5 |
| Speed | Trial 4 | 1.9±0.3 | 2.2±0.3 | 2.0±0.4 | 2.2±0.3 |
| Speed | Trial 5 | 1.9±0.2 | 2.2±0.4 | 1.9±0.4 | 2.2±0.4 |
| 90 DPI |  | WT sham | MAPTKI sham | CHI WT | MAPTKI CHI |
| Distance | Habituation | 27.0±3.2 | 28.4±2.6 | 29.6±2.9 | 34.7±7.2 |
| Distance | Acquisition Trial 1 | 13.2±2.0 | 16.2±2.2 | 12.4±1.2 | 15.4±2.6 |
| Distance | Trial 2 | 10.9±1.1 | 14.2±2.1 | 10.5±1.6 | 13.8±4.4 |
| Distance | Trial 3 | 10.2±1.1 | 13.6±2.9 | 10.9±1.1 | 14.6±3.7 |
| Distance | Trial 4 | 9.9±1.1 | 13.9±2.3 | 10.4±1.1 | 15.1±4.0 |
| Distance | Trial 5 | 9.9±0.8 | 13.4±2.6 | 10.6±1.7 | 15.2±4.8 |
| Speed | Habituation | 4.5±0.5 | 4.8±0.4 | 5.0±0.5 | 5.8±1.2 |
| Speed | Acquisition Trial 1 | 2.2±0.3 | 2.6±0.5 | 2.0±0.2 | 2.6±0.5 |
| Speed | Trial 2 | 1.8±0.2 | 2.2±0.5 | 1.7±0.3 | 2.3±0.7 |
| Speed | Trial 3 | 1.7±0.2 | 2.3±0.4 | 1.7±0.2 | 2.4±0.6 |
| Speed | Trial 4 | 1.7±0.2 | 2.3±0.4 | 1.7±0.2 | 2.5±0.7 |
| Speed | Trial 5 | 1.7±0.2 | 2.2±0.4 | 1.8±0.2 | 2.6±0.9 |

**Table 17.**
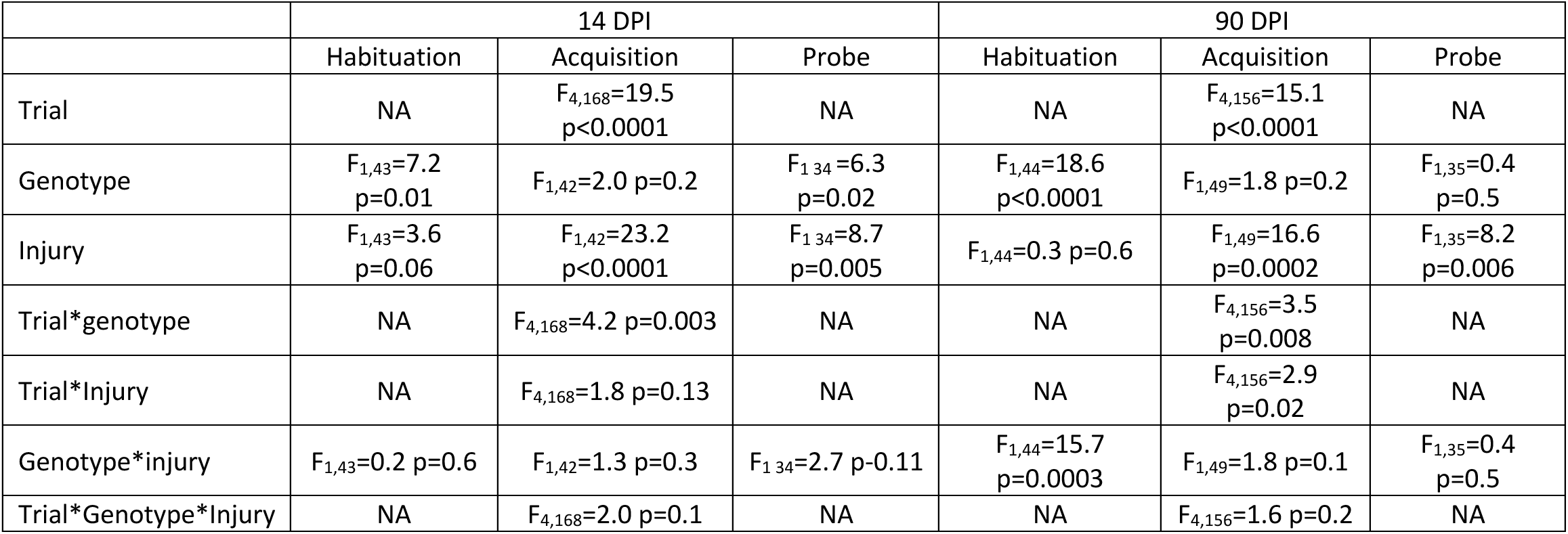
Statistical analysis of active place avoidance. Two-way ANOVA analyzes distance traveled in the habituation trial. Three-way ANOVA analyzes shock zone entrances in the acquisition trials. Two-way ANOVA analyzes shock zone entrances in the probe trial. NA, not analyzed.

During APA acquisition, shock zone entrances had significant effects of trial, injury, and trial*genotype but not genotype (Table 18). Injured MAPTKI mice learned the shock zone entrance while injured WT mice did not (Fig. 12A). WT or MAPTKI shams as well as injured MAPTKI mice significantly lower shock zone entrances from the first to fifth acquisition trials suggesting acquisition of the shock zone location (Fig. 12A). On the fifth and final acquisition trial, sham or injured MAPTKI mice have similar shock zone entries. In contrast, injured WT mice did not reduce shock zone entrances from the first to fifth trial (Fig. 12A).

**Figure 12.**
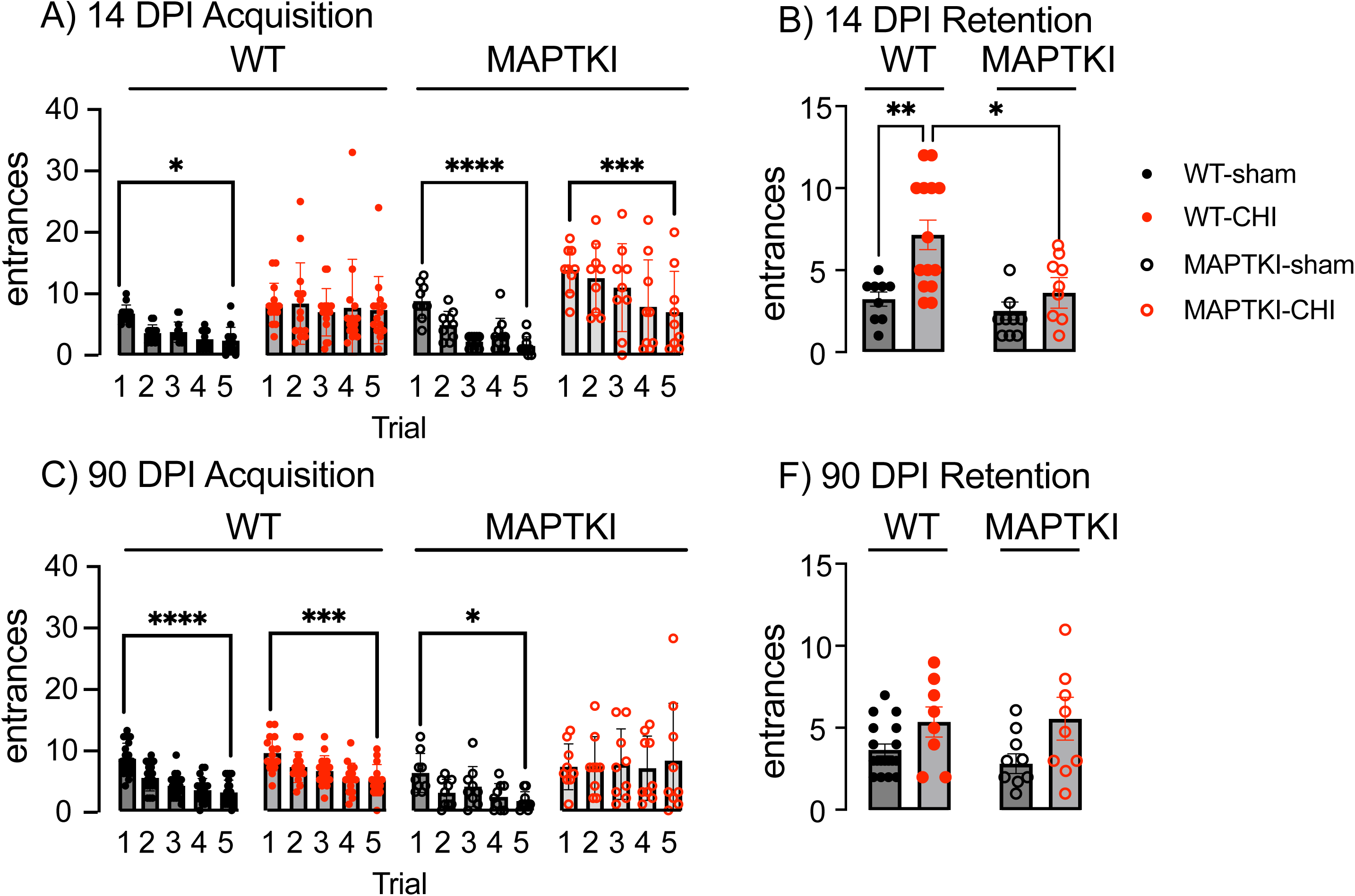
WT and MAPTKI mice differ in active place avoidance acquisition and retention. A) Summary of APA acquisition at 14 DPI. Training curves of WT (left) and MAPTKI (right) mice are graphed separately. Injured MAPTKI mice, but not WT mice lowered shock zone entrances between training trials 1 through 5. *p=0.02, ***p<0.0001**p=0.0002. B) Summary of APA retention at 14 DPI. *p=0.04, **p=0.004. C) Summary of APA Acquisition at 90DPI. Injured WT mice, but not MAPTKI mice lowered shock zone entrances between training trials 1 through 5. *p=0.02, **p=0.0002, ***p<0.0001. D) Summary of APA retention at 90 DPI. n’s=9-15

**Table 18.** Three-Way ANOVA analysis of alternating T-maze.

|  |  |
| --- | --- |
| Time | $F_{1,69}=3.5$ $p=0.07$ |
| Genotype | $F_{1,69}=7.5$ $p=0.008$ |
| Injury | $F_{1,69}=6.9$ $p=0.01$ |
| Time*genotype | $F_{1,69}=1.1$ $p=0.3$ |
| Time*injury | $F_{1,69}=4.5$ $p=0.04$ |
| Genotype*injury | $F_{1,69}=0.9$ $p=0.4$ |
| Time*genotype*injury | $F_{1,69}=4.7$ $p=0.03$ |

Three days after the final acquisition trial, a probe trial assesses retention of the shock zone location. Results from APA probe trials at 14 DPI from male and female mice are combined since sex did not affect APA entrances for either mouse strain (WT, F1,19=1.2 p=0.3; MAPTKI, F1,15=1.1 p=0.3). Entrances during probe trial have significant effects of genotype but not injury or injury*genotype (Table 18). On the probe trial, sham or injured MAPTKI mice show similar avoidance of the shock zone location, while injured WT mice entered the shock zone location more times than WT shams (Fig. 12B). These data suggest injured WT mice are impaired on APA while injured MAPTKI mice are not.

At 90DPI, separate cohort of WT and MAPTKI mice are assessed on APA acquisition and retention. Entrances have significant effects of trial, injury, and trial*genotype, but not genotype (Table 18). WT or MAPTKI shams as well as injured WT mice lowered the number of shock zone entrances between trials one and five. In contrast, injured MAPTKI mice did not lower shock zone entrances. These data suggest that at 90DPI, injured WT mice recover, whereas injured MAPTKI mice are now impaired. Results from APA probe trials at 90 DPI from male and female mice are combined since sex did not affect APA entrances for WT mice and, although significant, did not yield biologically significant comparisons for MAPTKI mice (WT, F_1,22_=3.7, p=0.07; MAPTKI, F_1,9_=5.4 p=0.04). All groups have similar entrances into the shock zone suggesting similar retention of the shock zone location. Taken together, these data suggest CHI induces a transient APA retention impairment in injured WT mice that resolves by 90 DPI. In contrast, injured MAPTKI mice develop impairments in both acquiring and retaining APA between 14 and 90 DPI. These data suggest differing cognitive deficits in injured WT and MAPTKI mice.

WT and MAPTKI mice are assessed on alternating T-maze at 14 and 90 DPI and the ability to alternate between the two T-maze arms is assessed using discrimination index. Results for male and females are combined since no significant sex effect is observed for WT and MAPTKI yields no significant biological pairwise comparisons (WT, F_1,50_=0.8 p=0.4; MAPTKI, F_1,12_=4.6 p=0.06). Discrimination index has significant effects of genotype, injury, time*injury, and time*genotype*injury (Table 18). At 14 DPI, discrimination index of injured WT is lower than shams (Fig.13). Discrimination index significantly lowers in sham WT mice between 14 and 90 DPI. Due to this age effect, discrimination index no longer differs between sham and injured WT mice (Fig. 13). These data suggest that, at 14 DPI, injury impairs working memory of WT, but not MAPTKI mice. At 90 DPI, an effect of age on sham WT mice obscures any injury effect on alternating T-maze.

**Figure 13.**
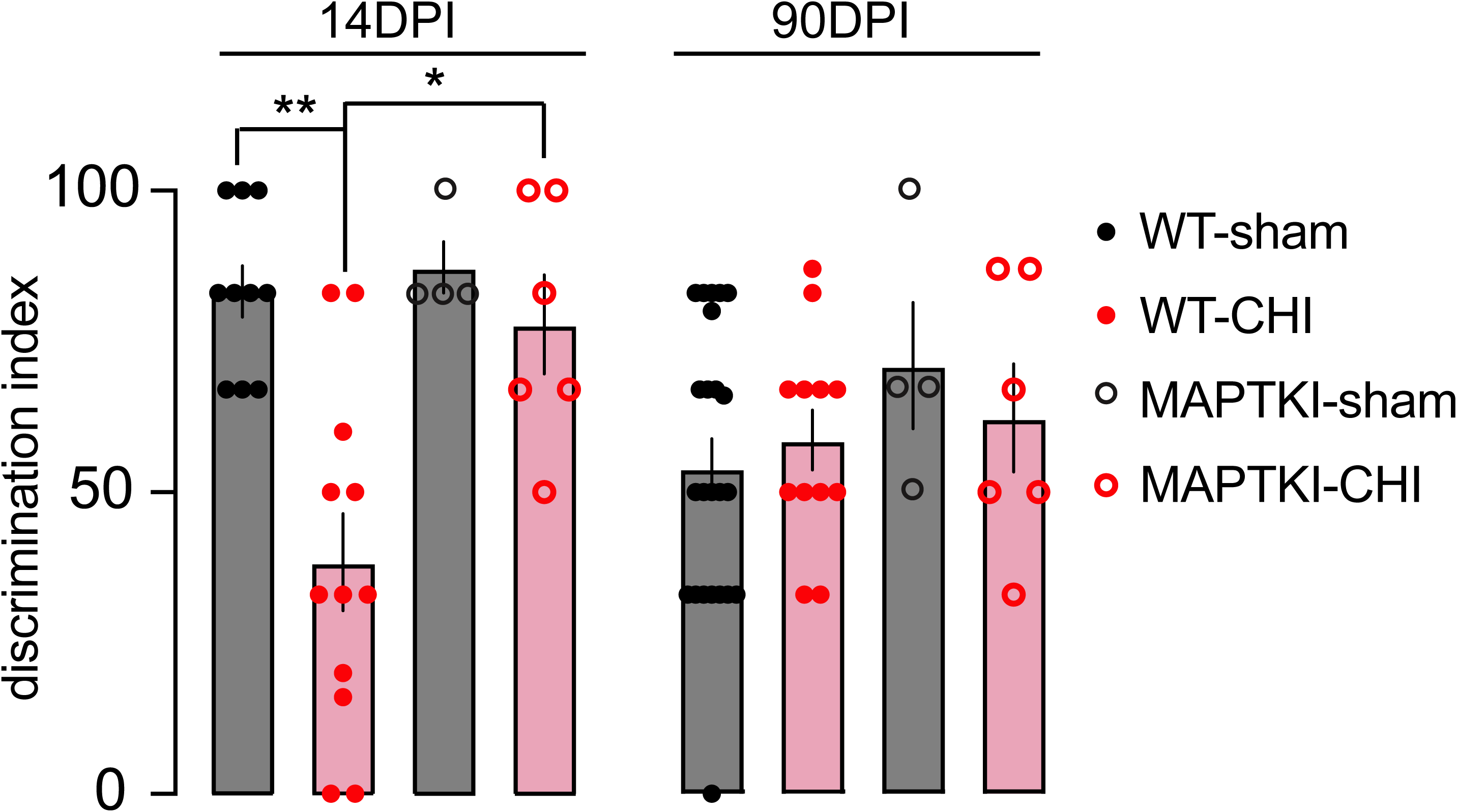
Injury impairs WT, but not MAPTKI mice, on alternating T-maze. At 14 DPI, discrimination index of injured WT mice is significantly lower than sham WT or injured MAPTKI mice. Discrimination index of di not differ among the sham and injured WT or MAPTKI groups. **p=0.002, *p=0.01. n’s=4-12.

## Discussion

This study demonstrates differing subacute and chronic disease courses in WT and MAPTKI mice. If left uninjured, WT or MAPTKI mice lack behavioral deficits, neurodegeneration, increased inflammation, or pTau accumulation up to 24 months of age [24, 32]. Thus, any differing disease trajectories of injured WT and MAPTKI mice must result from an interaction of genotype and injury and not from genotype alone. At the time of injury, WT and MAPTKI mice have similar body weight (Supplementary Figure 1A). WT and MAPTKI mice receive a similar initial injury as assessed by time to regain righting reflex (Supplementary Figure 1B). Similar initial injury is further suggested by similar gray matter injury to hippocampus, cortex and thalamus (Figures 1,2).

At 14 DPI, injured WT mice have greater subacute axotomy and overall axonal loss than MAPTKI mice (Figure 3). Axonal transport is impaired in WT, but not MAPTKI mice. The two strains, however, appears to have a similar loss of large caliper axons (Figure 5). At 14 DPI, injury produces myelin loss in WT, but not MAPTKI mice. By 90 DPI, corpus callosum myelin increases in WT mice content yet decreases in injured MAPTKI mice (Figure 6). Thus, the timing and extent of white matter damage differs substantially between the two strains. Injury also produced differing pTau expression in WT and MAPTKI mice (Figs. 7-9). At 90 DPI, injured WT and MAPTKI cortex have a similar density of pathological protein aggregates, yet the density of Thio-S+ aggregates is higher in MAPTKI corpus callosum and thalamus (Figure 10). At 90 DPI, the contralesional MAPTKI thalamus also has a delayed neuronal loss that is absent in WT mice (Figure 2). The high proportion of white matter in thalamus may explain why the MAPTKI thalamus has neuronal loss at 90 DPI [67, 68]. At chronic times, the corpus callosum of WT mice suggests remyelination while MAPTKI mice exhibit myelin loss. At 90 DPI, both WT and MAPTKI mice have Thio-S^+^ aggregates in cortex, yet only WT accumulates Thio-S+ aggregates in thalamus, while MAPTKI accumulates Thio-S+ aggregates in corpus callosum (Figure 10). Increased AT8 pTau cell density accompanies the elevated Thio-S^+^ cell density in WT and MAPTKI cortex or corpus callosum (Figures 7 and 10). These data suggest that despite similar initial injury and grey matter injury, MAPTKI mice have substantially less subacute white matter injury than WT. At chronic times white matter injury can worsen in MAPTKI, but not WT mice.

Behavioral testing also reveals differing disease courses in injured WT and MAPTKI mice. At 14 DPI, cognitive deficits in injured WT mice are substantial; they do not retain Barnes maze, neither acquire nor retain APA and are impaired in alternating T-maze (Figures 11-13). MATPKI both acquire and retain Barnes maze and APA and are not impaired on alternating T-maze. By 90 DPI, injured WT mice have 24-hour recall of Barnes maze and can now acquire APA while injured MAPTKI become impaired (Figure 11,12). These data suggest MAPTKI mice have fewer subacute behavioral deficits than WT. At chronic times, behavioral deficits in injured WT mice partially recover while deficits in injured MAPTKI worsen (Figures 12,13).

These data suggests that injured WT mice provide a model of the progression of TBI patients who ultimately recover from a subset of their subacute deficits. Multiple mouse TBI show a similar progression of recovery [69]. A subset of TBI patients, however, initially show fewer deficits, but ultimately get worse over time [70]. Few mouse models of TBI recapitulate this late onset of chronic injury [69, 71]. A single CHI to MAPTKI mice produces these late-onset, chronic deficits. These findings suggest injured MAPTKI provide a useful model to investigate disease progression following CHI that is more translatable to the subset of TBI patients developing later onset of chronic deficits.

Cognitive deficits differ between WT and MAPTKI mice. Both injured WT and MAPTKI mice have mild subacute impairments acquiring Barnes maze (Figure 11A). Whitney. et al. reported similar deficits in CHI-injured WT mice [20]. Despite similar acquisition, injured MAPTKI mice recall the escape hole location while WT mice do not (Figure 11B, C). Cognitive deficits have been reported in injured mice up to 2-month post-injury [11–14]. It remains uncertain if these injured mice improved at later times in a manner similar to injured WT CHI mice. By 90 DPI, WT and MAPTKI mice acquire Barnes maze, yet only MAPTKI have impaired recall (Figure 11C).

Acquisition of Barnes maze activates multiple connected brain structures including hippocampus, thalamus, and cortex [46]. APA depends upon the hippocampus and fimbria-fornix [47, 51]. At 14 DPI Injured WT and MAPTKI mice have similar neuronal loss in the ipsilesional hippocampus, cortex, and thalamus, yet injured WT mice have white matter damage absent in MAPTKI mice. This increased white matter damage may underlie the memory deficits of injured WT mice. (Figures 3,5 and 6). In contrast, at 90 DPI, white matter damage has increased in injured MAPTKI mice along with neuronal loss in the contralesional thalamus. This additional gray and white matter damage may contribute to the development of impaired memory in injured MAPTKI mice by 90 DPI.

APA deficits also differ between injured WT and MAPTKI mice. At 14DPI, injured MAPTKI mice acquire and retain the APA shock zone location while WT mice do not (Figure 12A,B). At 14DPI, injured WT and MAPTKI mice have similar gray matter injury but WT mice have greater white matter injury (Figures 3,5 and 6). White matter damage impairs APA acquisition [51, 72]. Increased white matter injury in injured WT mice likely underlies APA acquisition deficits at 14DPI. At 90DPI, injured WT mice acquire APA, whereas injured MAPTKI are impaired (Figure 12C). At 90DPI, injured WT mice acquire APA, despite maintaining similar cortical, hippocampal and thalamic neuronal loss present at 14DPI (Figures 1,2 12C). At 90 DPI, WT mice remyelinate the injured corpus callosum, in contrast to the demyelination of corpus callosum in injured MAPTKI (Figure 6). White matter repair may underlie the ability of WT, but not MAPTKI mice to acquire APA at 90 DPI.

At 14 DPI, Injured WT mice have alternating T-maze deficits that are absent in MAPTKI mice (Figure 13). Lesions to thalamus, hippocampus, or corpus callosum impair mice in alternating T-maze [53, 54]. At 14DPI, injured WT or MAPTKI have similar hippocampus and thalamus neuronal loss yet the injured WT corpus callosum has greater axonal damage and myelin loss (Figures 1-3,5,6). These data suggest that increased white matter injury in WT mice contribute to alternating T-maze deficits. These results also suggest that the similar neuronal loss in injured MAPTKI and WT mice at 14 DPI is insufficient to produce T-maze deficits. By 90DPI, sham groups alternate no better than injured groups suggesting an age-dependent decline in alternating T-maze. Others have described age-dependent deficits in alternating T-maze [53].

Less subacute white matter injury in MAPTKI mice suggests a protective effect of 3R tau. 3R tau has fewer microtubule binding sites and binds to microtubules with lower affinity than 4R tau [27]. *In vitro* studies suggest greater flexibility of 3R tau-containing axons [31]. While mechanical forces may bend and stretch both 3R and 4R containing axons, increased flexibility of 3R-containing axons may underlie the relative resistance of white matter in MAPTKI mice.

This study assessed both disease course in both male and female WT or MAPTKI mice. Males and female WT or MAPTKI mice had similar histological damage and cognitive deficits (Figures 1-6, 11-13). AT8 pTau expression did not differ between the sexes (Figure 7). PHF1 and S214 pTau expression are the only study outcomes that had a significant sex effect (Figures 8,9). Injury increases PHF^+^ pTau cell density at 14 DPI in the female MAPTKI cortex and the female WT thalamus (Figure 8B, C). In both cases, female MAPTKI and WT mice have a greater density of cells with the PHF1^+^ pTau phosphorylation site (Ser396, Ser 404) than males. Cortical S214 pTau expression is similar in both sexes. S214 pTau expression is higher in female WT corpus callosum and in male MAPTKI corpus callosum (Figure 9D, G). Few studies have reported sexual dimorphisms in tau phosphorylation in rodents. Aging or heat shock results in more PHF1 phosphorylation in female rats than males [73, 74]. Tauopathy is less severe in female mice expressing a pathogenic P301S mutant tau than males (Sun et al., 2020).

This study has multiple caveats. Chronic deficits of WT and MAPTKI mice are assessed only at 90 DPI. Previous studies showed chronic deficits in injured WT mice at 180 DPI [21]. Thus, the differing disease courses of WT and MAPTKI mice should be examined at times later than 90 DPI. Differing disease courses in WT and MAPTKI implies alterations in glial activation and the inflammatory response. These key aspects of brain injury remain to be assessed. In addition to differing expression of 3R and 4R tau, WT mice express murine tau while MAPTKI mice express human tau. Human and murine tau differ in their interaction with axonal, somatic, dendritic, and synaptic proteins [75]. The interactome of human tau with murine proteins in MAPTKI mice is unknown yet may contribute to the differing disease courses between injured WT and MAPTKI mice.

