## Supplementary figures and images for "Tau Isoform Expression Drives Disease Outcomes Following a Single Closed Head Injury"

### Supplemental figure 1.pdf

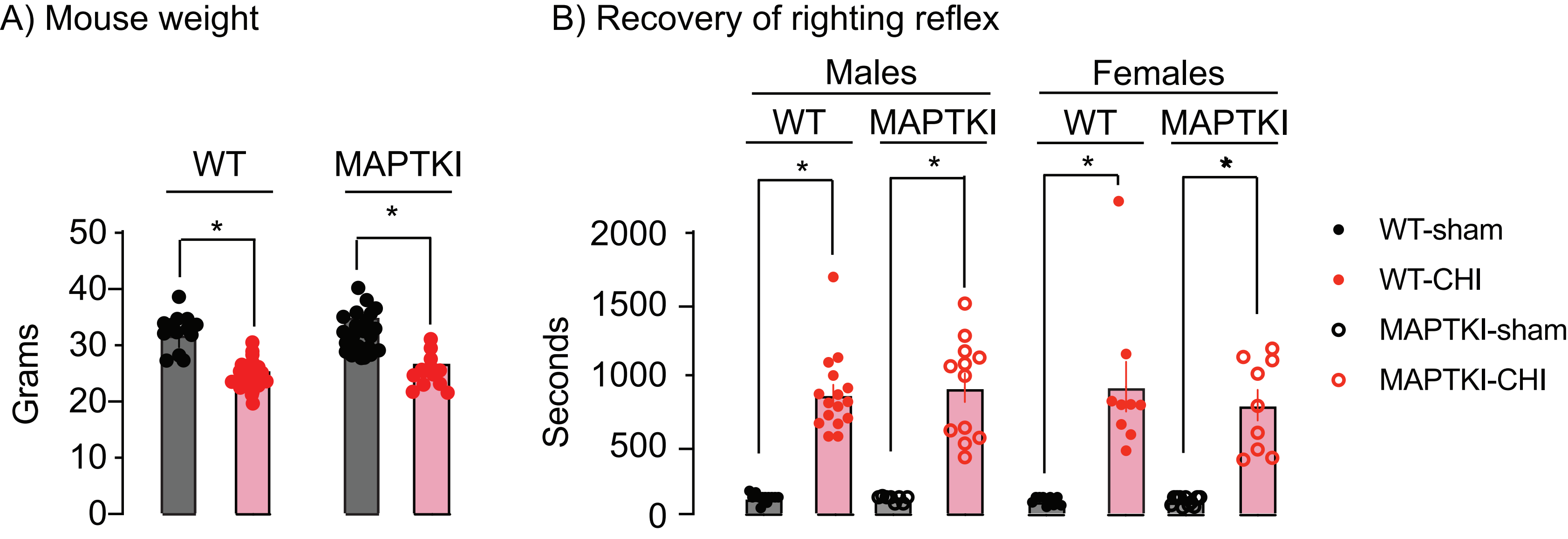
